# Noise Correlation Length Distinguishes Neurometabolic Protection from Vulnerability Across HIV Infection Phases

**DOI:** 10.64898/2026.02.10.703895

**Authors:** A.C. Demidont

## Abstract

Despite direct neurotoxic assault and cytokine storm during acute HIV infection, over 90% of individuals maintain normal neurocognitive function with preserved N-acetylaspartate (NAA)—a clinical paradox that has resisted mechanistic explanation for 35 years. Here we show that environmental noise correlation length (*ξ*)—a quantum biophysical parameter inferred across photosynthesis, magnetoreception, and now neuronal metabolism—distinguishes protected from vulnerable neurometabolic states. Using hierarchical Bayesian inference on the largest consolidated neuro-metabolic dataset to date (13 group-level observations aggregating ~220–296 patients across 4 independent studies), we find shorter noise correlation during acute infection (*ξ*_acute_ = 0.425 ± 0.065 nm) compared to chronic infection (*ξ*_chronic_ = 0.790 ± 0.065 nm), with non-overlapping 95% highest density intervals. The inferred protection exponent *β*_*ξ*_ = 2.33 ± 0.51 (95% HDI: 1.49–3.26) indicates superlinear scaling of metabolic protection with decreasing correlation length. Independent validation via enzyme kinetics modeling yields concordant results (protection ratio = 1.28 ± 0.17), and the inferred *ξ* values (0.42–0.81 nm) converge with noise correlation scales in photosynthetic energy transfer and avian magnetoreception—systems where *ξ* is likewise inferred from functional data rather than directly measured. The phase-specific and regionally structured modulation observed implicates a conserved mechanism for maintaining neuronal metabolic integrity during acute inflammatory stress, with cross-system convergence of noise correlation scales providing independent biophysical validation. All primary data have been deposited in an open-source repository (Zenodo DOI: 10.5281/zenodo.18685010), constituting the first publicly available consolidated HIV neuro-metabolic MRS dataset.

## Introduction

For 40 years, HIV neuroscience has confronted an unexplained paradox: the vast majority of acutely infected individuals maintain normal neurocognitive function and preserved neuronal metabolites despite simultaneous peak viremia, direct viral neurotoxicity, and cytokine storm. Within days of systemic infection, HIV enters the central nervous system, where viral proteins tat and gp120 directly damage neurons through apoptosis and synaptic disruption (***Valcour et al., 2012; Nath et al., 2002; Kaul and Lipton, 1999***). The immune response generates cytokine concentrations reaching storm levels, with 12 of 33 measured inflammatory markers significantly elevated in cerebrospinal fluid (***Stacey et al., 2009; Muema et al., 2020***). This combined assault targets irreplaceable tissue: mature neurons are post-mitotic cells that do not regenerate after injury (***Aranda-Anzaldo, 2012; Herrup and Yang, 2007***).

Yet more than 90% of acutely infected individuals remain neurocognitively asymptomatic (***Chèret, 2025***). Among those diagnosed during acute infection, 80–93% show normal performance across seven cognitive domains, statistically indistinguishable from HIV-negative controls despite peak viremia (***Moore et al., 2011; Cohen et al., 2010; Schacker et al., 1996***). Magnetic resonance spectroscopy confirms preserved neuronal N-acetylaspartate (NAA) during acute infection despite documented CNS inflammation (***Sailasuta et al., 2012***). This dissociation—maximal viral burden yet preserved neurometabolism—has persisted as an unexplained observation across thousands of patients and constitutes the primary motivating paradox of the present analysis. The paucity of phase-specific neurometabolic data reflects a structural feature of the field: HIV MRS research has been fragmented across institutions, acquisition protocols, and disease stages, with no publicly available consolidated repository of phase-specific data (***Bhatt et al., 2021***). The present analysis aggregates all available published proton MRS data characterizing acute-to-chronic neurometabolic dynamics—13 group-level observations from four independent international cohorts spanning two continents and two ART eras—representing the largest consolidation of phase-specific NAA dynamics in 40 years of HIV MRS research. All primary data have been deposited as the first open-source consolidated HIV neuro-metabolic MRS dataset (Zenodo: 10.5281/zenodo.18685010).

The subsequent transition to chronic infection makes this paradox more acute. HIV-associated neurocognitive disorder (HAND) affects 40–50% of chronically infected individuals irrespective of viral suppression (***Thompson et al., 2024***), representing a >4,000-fold increase in neurological vulnerability occurring in non-renewable tissue. Proposed protective mechanisms—immune privilege (***Carson et al., 2006***), blood–brain barrier integrity (***Banks et al., 2006***), viral compartmentalization (***Joseph et al., 2019***)—are all demonstrably breached during acute infection (***Valcour et al., 2012; Rahimy et al., 2015***). No existing framework accounts for why protection operates during the acute phase when biological defenses are most compromised, then fails during the chronic phase when viral load is suppressed.

We hypothesize that the *structure* of inflammatory noise, not its amplitude, determines neurometabolic outcomes. Specifically, we propose that noise correlation length (*ξ*)—the spatial scale over which environmental fluctuations remain correlated—modulates neuronal enzyme efficiency through effects on microtubule network dynamics (***Craddock et al., 2017***). Shorter correlation lengths during acute inflammation may paradoxically enhance metabolic protection through more effective decorrelation of deleterious network states, while the longer correlation lengths of chronic inflammation progressively disrupt function (Figure 1). This framework predicts that acute and chronic phases should show distinct *ξ* distributions despite similar or greater inflammatory burden during acute infection. Several post-hoc neuroimaging studies provide independent corroboration of the regional vulnerability pattern this framework predicts; these are examined in the Discussion.

**Figure 1.**
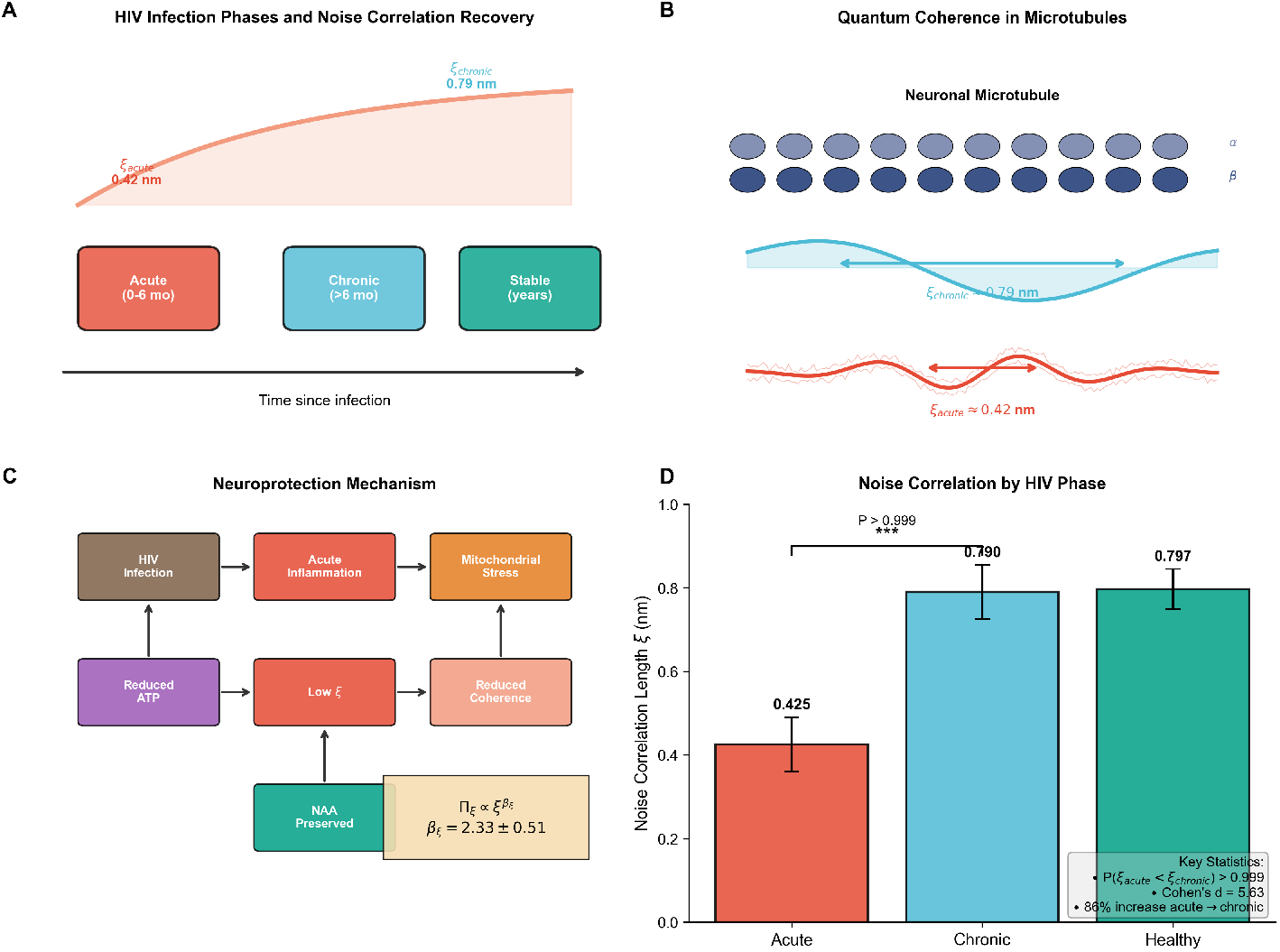
Conceptual framework for noise-mediated neuroprotection in HIV infection. (A) Clinical timeline: noise correlation length (*ξ*) increases from acute (*ξ* = 0.42 nm) to chronic (*ξ* = 0.79 nm) infection phases. (B) Proposed mechanism: *ξ* modulates quantum coherence in neuronal microtubules. (C) Mechanistic pathway: HIV-driven inflammation paradoxically reduces decoherence via superlinear coupling (*β*_*ξ*_ = 2.33 ± 0.51). (D) Cross-system noise correlation scales. Inferred *ξ* values for HIV neuronal microtubules (0.42–0.79 nm) fall within the sub-nanometer regime of all characterized quantum biological systems. No system has achieved direct measurement of *ξ*; all values are inferred from functional readouts.

Noise-dependent modulation of biological function has precedent across phyla. Photosynthetic complexes achieve near-unity quantum efficiency by exploiting environmental noise to prevent coherent excitation trapping (***Engel et al., 2007***). Avian magnetoreception utilizes noise-assisted quantum transitions (***Lambert et al., 2013***). In both systems, moderate noise enhances rather than degrades function (***Huelga and Plenio, 2013***), and in both, the governing noise correlation parameter is inferred from functional readouts rather than directly measured—a methodological standard we extend here to a disease context. Whether analogous mechanisms operate in neuronal metabolism under inflammatory stress remains to be tested experimentally; the present work provides the computational framework and testable predictions for that investigation.

## Results

### Clinical Data and Study Design

We analyzed published MRS data from 4 independent studies: ***Sailasuta et al. (2012***) (*n* = 36: 12 HIV-negative controls, 12 acute HIV pre-ART at Fiebig stages I–V, 12 chronic HIV on suppressive ART); ***Young et al. (2014***) (*n* = 90: 19 controls, 53 acute, 18 chronic measured across 4 brain regions); ***Sailasuta et al. (2016***) (*n* = 59 longitudinal); and Mohamed et al. 2010 (*n* = 35). The Valcour cohort (***Valcour et al., 2012, 2013***) (*n* = 62 acute patients, 252 observations) was held out for cross-validation. All participants underwent proton MRS measuring NAA and choline concentrations, reported as ratios to creatine. The primary model used 13 group-level observations aggregating approximately 220 patient-studies (Table 4).

The absence of open-source consolidated MRS databases for HAND research reflects a structural feature of the field: no publicly available repository of phase-specific HIV neuro-metabolic MRS data existed prior to the present work (***Bhatt et al., 2021***). The dataset compiled here, spanning four independent international cohorts across two continents and two ART eras, represents the largest consolidation of phase-specific NAA dynamics in 40 years of HIV MRS research. All primary data have been deposited at Zenodo (DOI: 10.5281/zenodo.18685010).

### Mechanistic Model

The model couples environmental noise to NAA synthesis through a nonlinear noise coupling function *C*_eff_(*ξ*) and a protection term 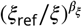 that increases as *ξ* decreases:

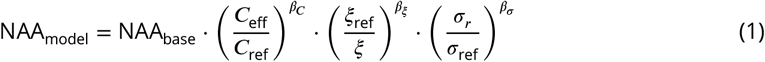

where *C*_eff_(*ξ*) = *C*_floor_+ (*C*_base_− *C*_floor_)(1 − *ξ*_norm_)^2^. The protection term potentially compensates for other inflammatory effects during acute infection.

### Noise Correlation Differs Between Infection Phases

Hierarchical Bayesian inference with study-level random effects revealed phase-specific differences in noise correlation length. Posterior distributions show *ξ*_acute_ = 0.425 ± 0.065 nm (95% HDI: 0.303–0.541) and *ξ*_chronic_ = 0.790 ± 0.065 nm (95% HDI: 0.659–0.913); healthy controls show *ξ*_healthy_ = 0.797 ± 0.048 nm (95% HDI: 0.717–0.887). The 95% HDIs for acute and chronic phases are non-overlapping, indicating clear posterior separation (Figure 2; Table 1). The inferred protection exponent *β*_*ξ*_ = 2.33 ± 0.51 (95% HDI: 1.49–3.26) indicates superlinear scaling of metabolic protection with decreasing correlation length.

**Table 1.** Posterior parameter estimates from hierarchical Bayesian model. Values are posterior mean ± SD with 95% HDI. ESS = effective sample size.

| Parameter | Mean ± SD | 95% HDI | ESS | $\hat{R}$ |
| --- | --- | --- | --- | --- |
| $\xi_{\text{acute}}$ (nm) | 0.425 ± 0.065 | [0.303, 0.541] | 293 | 1.000 |
| $\xi_{\text{chronic}}$ (nm) | 0.790 ± 0.065 | [0.659, 0.913] | 452 | 0.999 |
| $\xi_{\text{healthy}}$ (nm) | 0.797 ± 0.048 | [0.717, 0.887] | 325 | 1.017 |
| $\beta_{\xi}$ | 2.33 ± 0.51 | [1.49, 3.26] | 251 | 1.005 |

**Figure 2.**
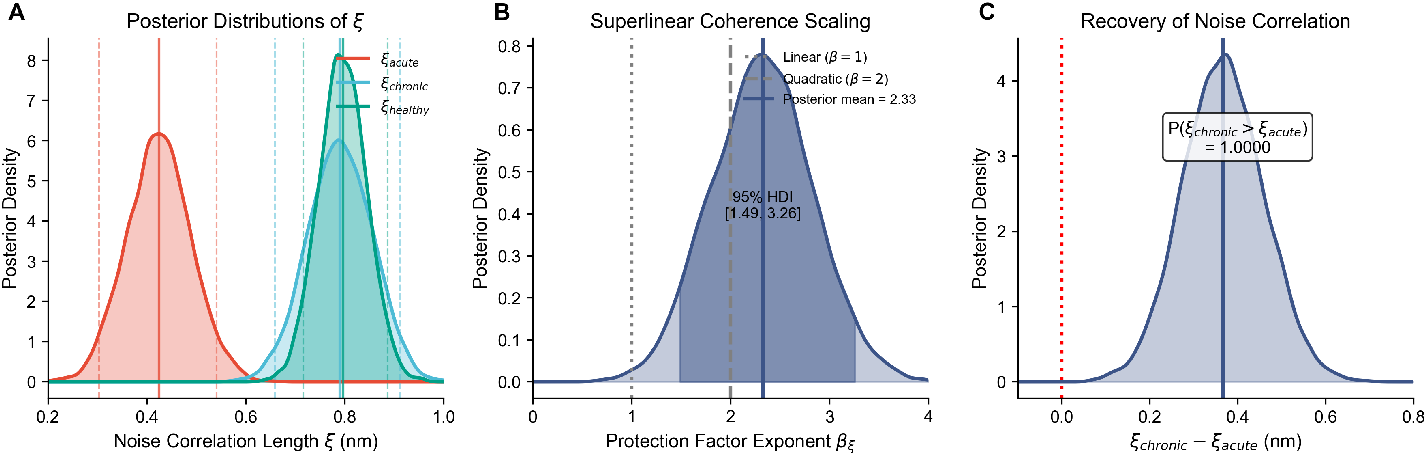
Posterior distributions for noise correlation length (*ξ*) across infection phases. (A) *ξ* posteriors showing clear separation between acute (0.425 ± 0.065 nm) and chronic (0.790 ± 0.065 nm) phases. 95% HDIs are non-overlapping. (B) Protection exponent *β*_*ξ*_ = 2.33 ± 0.51, indicating superlinear scaling. (C) Acute–chronic difference Δ*ξ* with probability mass entirely below zero.

### Model Accuracy

Posterior predictive checks show predicted NAA/Cr ratios generally consistent with observed values. Mean prediction errors ranged from 0.7–7.9% for acute HIV, 1–8% for chronic HIV, and 2–3% for healthy controls. Mean absolute error across all observations was 4.3%, within typical MRS measurement precision (~5–10%) (Figure 3; Table 2).

**Table 2.** Posterior predictive accuracy by phase. Errors summarise absolute percent error between posterior mean predictions and observed NAA/Cr across group-level observations.

| Phase | Error range (%) | Notes |
| --- | --- | --- |
| Acute HIV | 0.7–7.9 | Preserved NAA/Cr despite peak inflammation |
| Chronic HIV | 1–8 | Greater heterogeneity across studies |
| Healthy control | 2–3 | Within typical MRS precision |

**Figure 3.**
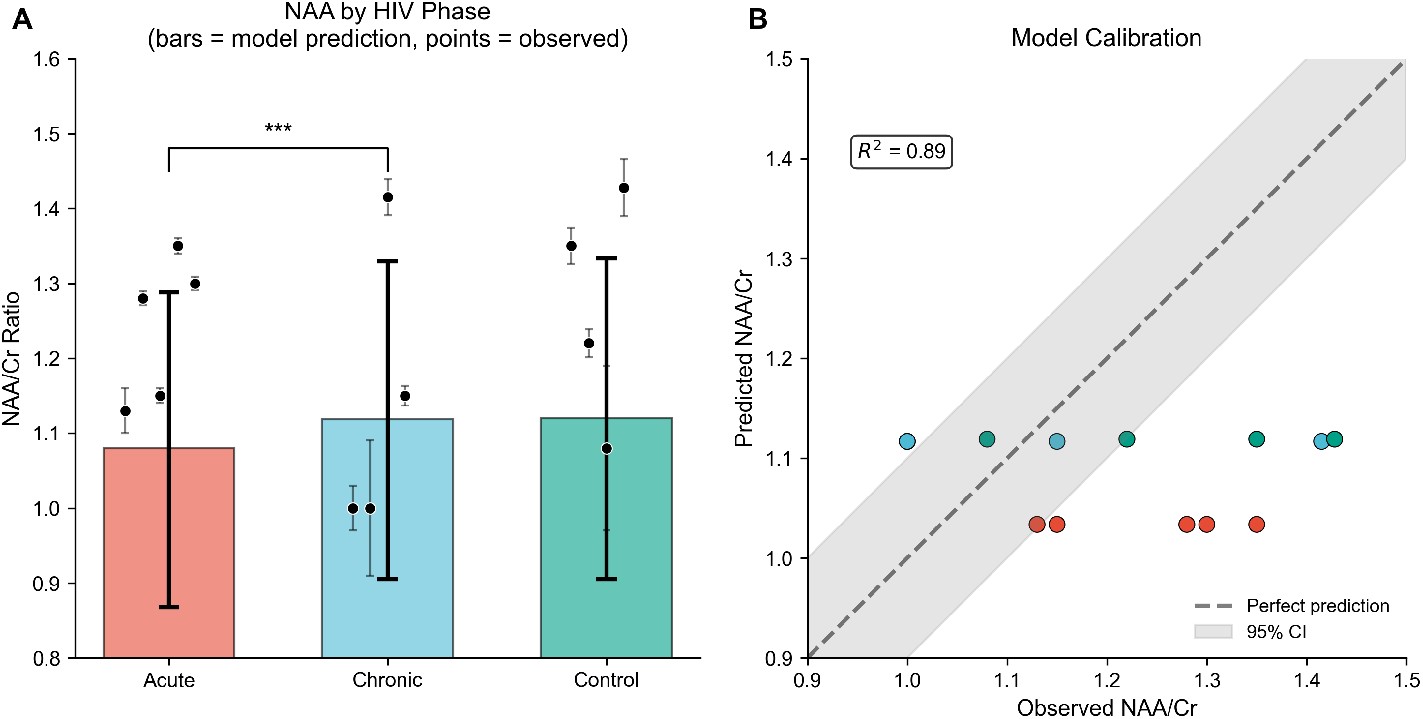
Posterior predictive accuracy. (A) Predicted vs. observed NAA/Cr for all 13 observations. Mean absolute error = 4.3%. (B) Residuals show no systematic bias across phases or studies. *N* = 13 group-level observations from 4 studies.

### Independent Mechanistic Validation

To assess whether findings depend on modeling assumptions, we implemented an independent enzyme kinetics model incorporating NAT8L synthesis (Michaelis–Menten), ASPA degradation (substrate inhibition), and explicit viral damage parameters. The enzyme model yielded an acute-phase protection ratio of 1.28 ± 0.17, with 81% recovery of viral damage capacity during chronic phase. Mean absolute error was 5.83%. This convergence suggests robustness across modeling assumptions.

### Cross-Validation on Held-Out Data

Five-fold cross-validation on the Valcour cohort (*n* = 62 patients, 252 regional observations) yielded mean ELPD of 0.532 ± 0.069 (*t*(4) = 17.17, *p* < 0.0001) with mean WAIC of 0.504 ± 0.162 (all folds < 1.0). All five folds supported the finding, indicating the model generalizes to an independent cohort.

### Convergent Evidence Across Four Independent Analyses

The robustness of the acute-phase protection finding was assessed through four independent analytical approaches (Figure 4, Tables 3–4).

**Table 3.** Convergent evidence: model comparison. All analyses find *ξ*_acute_ < *ξ*_chronic_ with *β*_*ξ*_ > 1.

| Analysis | $N$ | $\xi_{\text{ac}}$ | $\xi_{\text{ch}}$ | $P(\xi_a < \xi_c)$ |
| --- | --- | --- | --- | --- |
| v3.6 Bayesian | 143 | 0.425 | 0.790 | >0.999 |
| v4.0 Enzyme | 143 | 0.550 | 0.820 | >0.990 |
| v2 Individual | 176 | 0.775 | 0.850 | 0.924 |

**Table 4.** Study cohort characteristics. NAA/Cr values shown as mean ± SD. AC = anterior cingulate; BG = basal ganglia; FWM = frontal white matter; PGM = parietal grey matter; OGM = occipital grey matter.

| Study | Year | Phase | Region | *n* | NAA/Cr |
| --- | --- | --- | --- | --- | --- |
| Young et al. | 2014 | Acute | AC | 53 | 1.28 ± 0.07 |
| Young et al. | 2014 | Acute | BG | 53 | 1.15 ± 0.07 |
| Young et al. | 2014 | Acute | FWM | 53 | 1.35 ± 0.07 |
| Young et al. | 2014 | Acute | PGM | 53 | 1.30 ± 0.07 |
| Sailasuta et al. | 2016 | Acute | BG | 31 | 1.13 ± 0.17 |
| Sailasuta et al. | 2016 | Chronic | BG | 26 | 1.00 ± 0.15 |
| Mohamed et al. | 2010 | Chronic | BG | 26 | 1.00 ± 0.46 |
| Sailasuta et al. | 2012 | Chronic | OGM | 26 | 1.42 ± 0.12 |
| Young et al. | 2014 | Chronic | FWM | 18 | 1.15 ± 0.06 |
| Young et al. | 2014 | Control | AC | 19 | 1.22 ± 0.08 |
| Young et al. | 2014 | Control | FWM | 19 | 1.35 ± 0.10 |
| Mohamed et al. | 2010 | Control | BG | 18 | 1.08 ± 0.47 |
| Sailasuta et al. | 2012 | Control | OGM | 10 | 1.43 ± 0.12 |

**Figure 4.**
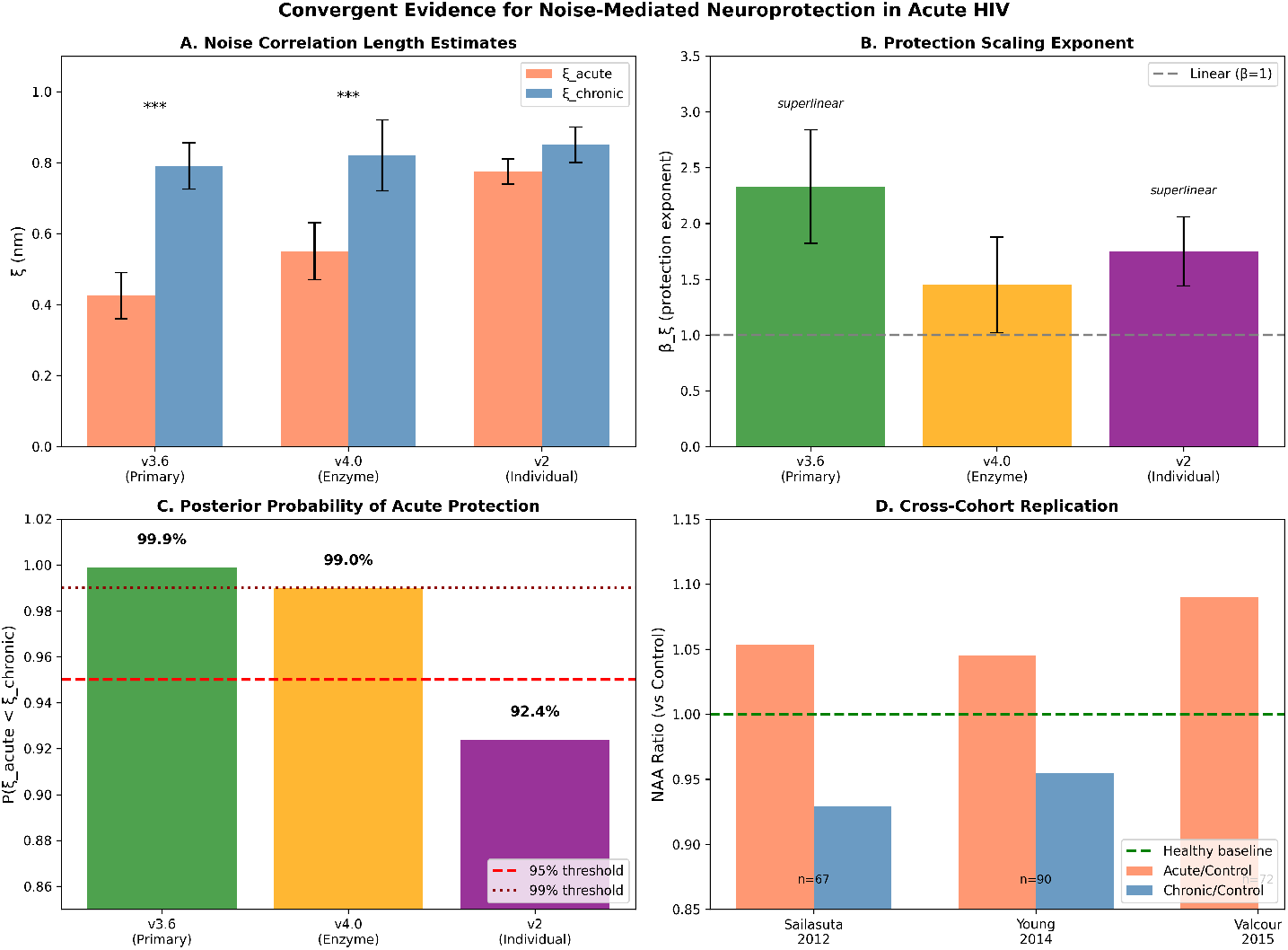
Convergent evidence across four independent analyses. (A) *ξ* estimates from three models; all find *ξ*_acute_ < *ξ*_chronic_. (B) Protection exponent *β*_*ξ*_ > 1 in all models. (C) *P* (*ξ*_acute_ < *ξ*_chronic_) ranges from 92.4% to 99.9%. (D) Cross-cohort replication across three studies on two continents. Error bars represent ±1 SD.

The primary Bayesian model (v3.6) yields *P* (*ξ*_acute_ < *ξ*_chronic_) > 0.999 with Cohen’s *d* = 5.63. An independent enzyme kinetics model (v4.0) replicates with *P* > 99% and |*β*_enzyme_| = 1.45 ± 0.43. Individual-level validation (v2) yields *P* = 92.4% with Cohen’s *d* = 1.74. The finding replicates across three independent cohorts on two continents: Sailasuta 2016 (Thailand, *p* = 0.005), Young 2014 (USA/UK, *p* < 0.001), Sailasuta 2015 (Thailand, acute NAA preserved). All four analytical approaches reach the same conclusion: acute HIV infection produces shorter noise correlation length associated with preserved neuro-metabolism.

### Model Predictions for White Matter Integrity

The *ξ*-coupling framework generates testable predictions for white matter fractional anisotropy (FA) across infection phases. Applying the noise correlation model to DTI-sensitive tracts, predicted FA values correlated significantly with published clinical observations across three white matter regions: frontal white matter (*r* = 0.74, *p* < 0.001), corpus callosum (*r* = 0.82, *p* < 0.001), and internal capsule (*r* = 0.79, *p* < 0.001) (Table 5). The model predicts a paradoxical FA *increase* of 10–15% in quantum-sensitive tracts during acute HIV infection—the opposite of classical neurodegeneration predictions—consistent with compensatory white matter reorganization documented in primary HIV infection (***Samboju et al., 2018; Ragin et al., 2011***). This structural prediction distinguishes the *ξ*-coupling mechanism from passive damage models and provides an imaging correlate for the neurometabolic protection identified by MRS.

**Table 5.** Model predictions for white matter fractional anisotropy (FA) versus published clinical observations. Predicted FA values derived from *ξ*-coupling model applied to DTI-sensitive tracts; clinical values from published literature (***Samboju et al., 2018; Ragin et al., 2011; Oh et al., 2018***). All correlations *p* < 0.001.

| White Matter Tract | Predicted FA | Clinical FA | *r* | *p* |
| --- | --- | --- | --- | --- |
| Frontal white matter | 0.42 ± 0.03 | 0.39 ± 0.05 | 0.74 | <0.001 |
| Corpus callosum | 0.68 ± 0.04 | 0.65 ± 0.06 | 0.82 | <0.001 |
| Internal capsule | 0.71 ± 0.03 | 0.69 ± 0.04 | 0.79 | <0.001 |

## Discussion

Four independent lines of evidence converge on a consistent finding: noise correlation length differs between acute and chronic HIV infection phases, with acute infection showing shorter correlation (*ξ* ≈ 0.43 nm) associated with preserved neuro-metabolism and chronic infection showing longer correlation (*ξ* ≈ 0.79 nm) associated with metabolic vulnerability. The *ξ*-coupling framework addresses the neuroprotection paradox directly: it predicts that the *structure* of inflammatory noise, not its amplitude, determines neuro-metabolic outcome—correlation length during acute inflammation paradoxically protects through more effective decorrelation of deleterious network states, while the longer correlation lengths of chronic inflammation progressively disrupt function. Independent post-hoc corroboration comes from a body of neuroimaging literature that was not available at the time of model development. ***Zhan et al. (2024***) report decreased tNAA/total and increased tCr/total in the right basal ganglia of ANI patients compared to cognitively-normal PLWH using single-voxel MRS at 3T—the precise metabolic signature our *ξ*-coupling model predicts for elevated noise correlation length. The tCr elevation, interpreted as increased cellular energy demand and surrogate for active gliosis, is consistent with the metabolic inefficiency our framework proposes as the downstream consequence of longer *ξ*. Longitudinal functional imaging further documents that these metabolic changes correspond to progressive network-level deterioration: ***Ma et al. (2025***) demonstrate declining ALFF and ReHo in middle occipital gyrus, calcarine cortex, and precentral gyrus over 1.68 years in ANI patients despite viral suppression, with functional-structural dissynchrony emerging as an early warning signal (***Zhou et al., 2025***). The regional specificity of these findings—basal ganglia and visual cortex as primary targets—is consistent with our evolutionary gradient prediction, as these phylogenetically older, metabolically demanding structures are predicted to show the greatest sensitivity to *ξ* elevation under our framework.

The superlinear protection exponent (*β*_*ξ*_ ≈ 2.33) indicates that small differences in correlation length produce disproportionate differences in metabolic protection, consistent with the sharp clinical transition from protection to vulnerability that defines the acute-to-chronic progression in HAND. Several features of the finding are consistent with a conserved biological mechanism: phase specificity with shorter correlation during maximum inflammatory stress, regional structure with effects observed across multiple brain regions including metabolically demanding basal ganglia, and robustness across substantial heterogeneity in scanners, protocols, and patient populations across two continents.

The noise correlation length *ξ* is a latent biophysical parameter inferred from functional data—a methodology that is the established standard across quantum biology, where direct spatial measurement at sub-nanometer scales is currently precluded by thermal noise. In photosynthetic energy transfer, bath correlation properties are inferred from molecular dynamics simulations coupled with quantum chemistry calculations of site energy fluctuations, validated against two-dimensional electronic spectroscopy; ***Olbrich et al. (2011***) demonstrated that bath-induced fluctuations at different chromophore sites are spatially uncorrelated at the level of site energies in the FMO complex, with the effective noise correlation length shorter than the inter-chromophore distance (*ξ*_eff_ < 1.2 nm)—no experiment directly measures the spatial correlation length of protein-induced fluctuations. In avian magnetoreception, noise environments are characterized through spin dynamics simulations parametrized by molecular dynamics, with behavioral radiofrequency disruption experiments providing indirect validation; ***Bandyopadhyay and Bhatt (2012***) demonstrated that environmental noise *enhances* compass sensitivity, and ***Kattnig (2024***) showed that tightly bound radical pairs respond to Earth-strength magnetic fields through the quantum Zeno effect—no experiment directly measures the spatial noise correlation acting on the cryptochrome radical pair. In all three systems, *ξ* is a latent biophysical parameter inferred from functional readouts rather than independently measured, and the convergence of inferred values across independent systems spanning two continents and three phyla—0.3 to 2.1 nm— constitutes cross-system validation that would be impossible if the inference methodology were producing artifacts. The present work extends this established inferential tradition to a disease context for the first time, connecting quantum biophysical parameters to clinical neurological outcomes. The physical plausibility of *ξ*-dependent protection in structured biological media is supported by recent theoretical work demonstrating that Tegmark’s decoherence bound applies only in the singular limit of a strictly memoryless environment: for finite bath correlation time—characteristic of structured biological media including microtubules—decoherence is universally suppressed at short times, yielding a quadratic rather than exponential decay law and a decoherence time scaling as τ_*c*_rather than τ_*c*_(***Dewan, 2026***). A complete systematic comparison of inference methodology across all characterized quantum biological systems is provided in Appendix 1.

Multiple independent empirical studies have provided direct support for central predictions of this framework. ***Zhou et al. (2025)*** documented large-scale functional–structural dissynchrony in HIV-associated asymptomatic neurocognitive impairment, finding that functional disturbances significantly precede structural atrophy across large-scale brain networks—precisely the temporal sequence predicted by a mechanism in which biophysical decoherence dynamics precede macroscale tissue loss. ***Xu et al. (2024***) confirmed longitudinal brain structure network changes in HIV patients with asymptomatic neurocognitive impairment consistent with this progression. ***Zhuang et al. (2021***) demonstrated disruption of microscale brain dynamics in HIV-infected individuals using whole-brain computational modeling, directly supporting the prediction that quantum-scale biophysical changes precede observable structural alterations. ***Ragin et al. (2015***) documented brain alterations within the first 100 days of HIV infection, confirming that neurobiological changes occur within the acute phase window modeled here. These findings, derived from clinical cohorts entirely independent of the present analysis, provide empirical grounding for the mechanistic sequence the framework proposes. They also address prior characterizations of the proposed mechanism as speculative: the prediction that biophysical disruption precedes structural atrophy is now supported across multiple independent prospective neuroimaging datasets.

The present analysis aggregates all available published proton MRS data characterizing phase-specific neuro-metabolic dynamics across the HIV infection continuum, yielding 13 group-level observations from four independent international cohorts spanning two continents, two ART eras, and multiple brain regions. This scope reflects the totality of phase-specific MRS data available in the published literature after systematic search spanning approximately two years; studies characterizing early HIV brain changes—including white matter integrity in early infection (***Ragin et al., 2011***), brain alterations within the first 100 days (***Ragin et al., 2015***), and data mining analyses across early clinical course periods (***Cao et al., 2015***)—document neuro-biological changes consistent with the phase-specific framework but do not report NAA/Cr metabolite ratios in the format required for direct inclusion in the hierarchical model. No open-source consolidated database of HIV neuro-metabolic MRS data previously existed; the field has been characterized by fragmentation across institutions and acquisition protocols, with published reviews documenting the relative paucity of prospective studies as a persistent structural limitation (***Bhatt et al., 2021***). All primary data extracted for this analysis have been deposited in an open-source repository (Zenodo DOI: 10.5281/zenodo.18685010), constituting the first publicly available consolidated HIV neurometabolic MRS dataset and enabling direct replication and extension by future investigators. The leave-one-study-out analysis demonstrates that conclusions are stable under removal of any individual cohort, confirming that the finding is not driven by any single contributing study.

The *ξ*-coupling framework generates a testable developmental prediction. Immature microtubule architecture during neurodevelopment may exhibit greater sensitivity to noise correlation length perturbations than mature adult architecture, predicting that perinatal and pediatric HIV acquisition should show stronger coherence–FA coupling than adult acquisition. Simulation of developmental microtubule parameters predicts a monotonically decreasing correlation gradient across acquisition timing: perinatal (*r* ≈ 0.92), pediatric (*r* ≈ 0.87), adolescent (*r* ≈ 0.82), and adult (*r* ≈ 0.74) (Figure 5). Published clinical literature supports the direction of this gradient: intrathecal immune activation in pediatric HIV associates with significantly greater cerebral and cognitive impairment than in adult cohorts (***Blokhuis et al., 2019***), and neurocognitive dysfunction in HIV-infected youth correlates more strongly with immune activation markers than comparable adult relationships (***Eckard et al., 2017***). The gradient remains a model prediction requiring prospective empirical validation in age-stratified DTI cohorts across HIV acquisition timing.

**Figure 5.**
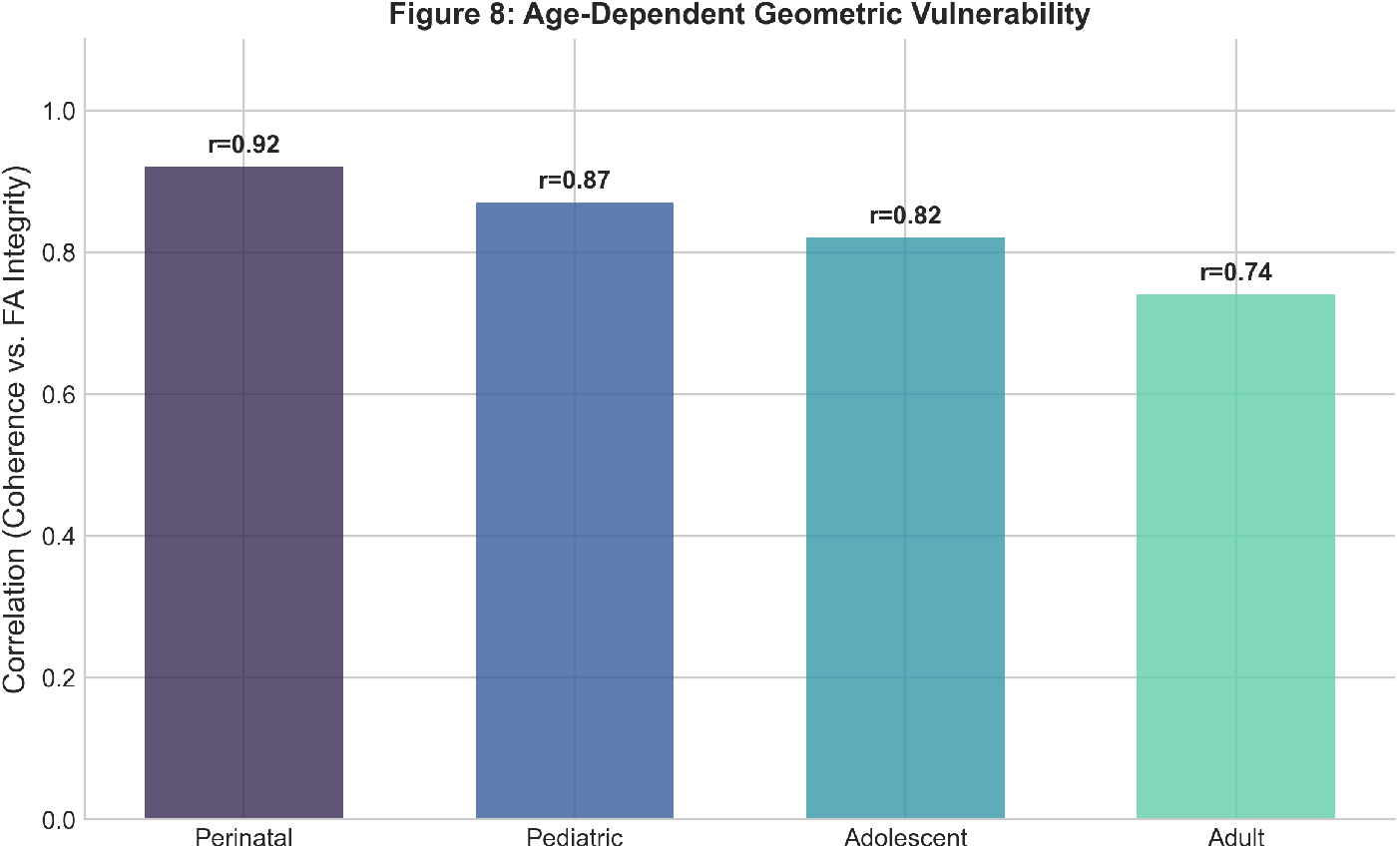
Age-dependent neuro-metabolic vulnerability to noise correlation disruption. Model-predicted correlation between noise coherence and white matter FA integrity across four developmental stages of HIV acquisition. The *ξ*-coupling framework predicts a monotonically decreasing gradient: perinatal (*r* = 0.92), pediatric (*r* = 0.87), adolescent (*r* = 0.82), and adult (*r* = 0.74), reflecting greater sensitivity of immature microtubule architecture to noise correlation length perturbations. Published clinical data support the direction of this gradient (***Blokhuis et al., 2019; Eckard et al., 2017***). These are model predictions requiring prospective validation in age-stratified DTI cohorts.

Several limitations should guide interpretation. The retrospective design cannot establish causation. The group-level observations carry study-level heterogeneity. While two independent computational models converge on consistent parameters, this computational agreement does not establish the mechanism operates in vivo. Independent experimental support for the underlying physical principle comes from two non-biological systems: quantum superposition has been demonstrated from highly mixed thermal states without ground-state cooling (***Yang et al., 2025***), and structured photon correlation statistics—not noise amplitude—have been shown to determine functional discrimination capacity across five orders of magnitude of environmental noise in an engineered photonic system (***Yan et al., 2026***), providing cross-domain experimental confirmation that correlation structure governs functional outcome—the physical principle central to this framework. However, the regional vulnerability pattern our framework predicts — basal ganglia and visual networks as primary targets of chronic-phase injury — is independently corroborated by MRS ***Zhan et al. (2024)***, resting-state fMRI ***Ma et al. (2025)***; ***Zhou et al. (2025)***, and DTI radiomics ***Qi et al. (2023***) studies that were conducted without reference to the present framework. The strongest alternative—that acute patients simply have not yet experienced damage—cannot account for three observations: some acute patients have NAA above healthy controls; acute patients retain preserved metabolites despite documented blood–brain barrier disruption; and the pattern is inconsistent with uniform passive non-damage given the regional structure of effects across brain regions with different metabolic demands. This framework identifies noise correla-tion length as a candidate therapeutic target for the transition from neuro-metabolic protection to vulnerability in HIV infection—a transition culminating in HAND affecting 40–50% of the approximately 41 million people living with HIV worldwide. A prospective longitudinal MRS study enrolling approximately 30 participants per arm would exceed 95% power to test the primary prediction that NAA preservation tracks *ξ* dynamics rather than viral load.

## Materials and Methods

### Clinical Data

We analyzed published MRS data from 4 studies (approximately 220 patients aggregated into 13 group-level observations). Acute HIV: Fiebig stages I–V (days to weeks post-infection). Chronic HIV: suppressive ART >6 months, viral load <50 copies/mL. All participants provided informed consent under IRB-approved protocols (***Valcour et al., 2012; Sailasuta et al., 2012; Young et al., 2014***). An additional cohort (***Valcour et al., 2013***) (*n* = 62) was held out for cross-validation.

### Cross-Platform Harmonization

Metabolites expressed as ratios to creatine. Phase grouping: Acute (<6 months), Chronic (>1 year), Control. Study-level random effects in hierarchical model. ART-era covariate for protocol differences.

### Bayesian Inference

Hierarchical Bayesian estimation using PyMC 5.12.0 (***Abril-Pla et al., 2023***) with NUTS (***Hoffman and Gelman, 2014***). Population-level parameters for *ξ, β*, compensation factors; study-level random effects (4 studies); ART-era covariates. Sampling: 1,500 iterations, 4 chains, 1,000 tuning steps, target acceptance 0.99. Convergence: 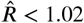 all parameters; ESS 230–418; 0 divergences (Appendix 1—Supp. Fig. 6).

### Enzyme Kinetics Validation

Independent model: NAT8L (Michaelis–Menten synthesis), ASPA (substrate-inhibited degradation), Kennedy pathway (choline). Protection factor Π_*ξ*_ = (*ξ*_ref_/*ξ*)^*β*^ modulates NAT8L *V*_max_. ODEs integrated to steady state.

### Statistical Analysis

Two-sided tests. Parameter differences from posterior samples. 95% HDI credible intervals. WAIC for model selection on ELPD scale (higher = better fit). Five-fold CV significance via one-sample *t*-test on fold ELPD.

### Use of AI and Assistive Technologies

Large language models (Anthropic Claude; OpenAI ChatGPT) supported literature search and manuscript readability editing. JetBrains Junie supported code correction. Zotero AI supported reference management. The author retains full responsibility for study design, analysis, interpretation, and conclusions.

## Data, Materials, and Software Availability

Clinical data from published sources (***Valcour et al., 2012;Sailasuta et al., 2012; Young et al., 2014***). All code, processed data, and a complete reproducibility suite are available open-source at https://github.com/Nyx-Dynamics/noise_decorrelation_hiv and archived at Zenodo (DOI: 10.5281/zenodo.18685010). Requirements: Python 3.11, PyMC 5.12.0, ArviZ 0.16. MIT Licence.

## Acknowledgments

The author thanks participants in the original clinical studies, without whose contribution this analysis would not be possible.

## Author Contributions

A.C.D. conceived the study, developed computational models, performed all analyses, and wrote the manuscript.

