## Supplemental Information, Tables and Figures for "Noise Correlation Length Distinguishes Neurometabolic Protection from Vulnerability Across HIV Infection Phases"

### 1 Cross-System Validation of Noise Correlation Scales in Quantum Biology

#### 1.1 Motivation

The noise correlation length parameter $\xi$ is inferred from metabolite data via hierarchical Bayesian modeling. Here we place this inference in the broader context of quantum biology, demonstrating that **no direct measurement of spatial noise correlation length exists in any quantum biological system**. Our inference of $\xi$ from clinical MRS data parallels—and extends—the approaches used in the two most established quantum biology systems: photosynthetic energy transfer and avian magnetoreception.

#### 1.2 Three Quantum Biological Systems: A Systematic Comparison

##### 1.2.1 Photosynthetic Energy Transfer (FMO Complex)

The Fenna–Matthews–Olson (FMO) pigment-protein complex in green sulfur bacteria achieves near-unity quantum efficiency in excitation energy transfer (EET). Seven bacteriochlorophyll *a* (BChl *a*) chromophores are arranged within a protein scaffold at average nearest-neighbor distances of ∼12 Å (1.2 nm) [10]. The bath correlation structure of the protein environment has been characterized by molecular dynamics simulations coupled with semiempirical quantum chemistry (ZINDO/S) calculations of site energy fluctuations [11].

Critically, Olbrich *et al.* found that bath-induced fluctuations at different chromophore sites are spatially **uncorrelated** at the level of site energies in the FMO complex [11]. This means the effective noise correlation length is shorter than the inter-chromophore distance ($\xi_{\text{eff}} < 1.2$ nm). Theoretical studies confirm that when spatial correlations are imposed with ∼30 Å decay length, exciton transport rates decrease approximately threefold [1]. The protein scaffold has evolved to maintain short bath correlations, precisely the regime our model identifies as neuroprotective.

Recent work using numerically exact non-perturbative simulations demonstrated that the detailed structure of the phonon spectral density—not a coarse-grained approximation—is critical for predicting coherence lifetimes, with excitonic coherences persisting on picosecond timescales at physiological temperature [8]. Environmental noise structure determines functional outcomes.

**Key parallel to this work**: Short bath correlation length ($< 1.2$ nm) enables efficient quantum transport. No direct measurement of $\xi$ was performed; the correlation structure was *inferred* from MD simulations and validated against 2D electronic spectroscopy data.

##### 1.2.2 Avian Magnetoreception (Cryptochrome 4a)

Migratory birds navigate using the Earth's magnetic field (∼50 $\mu$T) via cryptochrome flavoproteins in the retina. In European robin cryptochrome 4a (ErCry4a), blue-light excitation generates a series of radical pairs along a tryptophan tetrad [12, 13]. The magnetically sensitive radical pair RPC ($\text{FAD}^{\bullet-}$—$\text{TrpC}^{\bullet+}$) has an inter-radical separation of ∼17.6 Å (1.76 nm), while RPD has a separation of ∼21.3 Å (2.13 nm) [4].

A remarkable finding, directly paralleling our work, is that environmental noise can **enhance** compass sensitivity. Bandyopadhyay and Bhatt demonstrated that the sensitivity of the avian compass is enhanced by environmental noise, and that long coherence time is not required for navigation and may even spoil it [3]. Similarly, Kattnig showed that tightly bound radical pairs—previously thought insensitive to weak fields—can respond to Earth-strength magnetic fields through asymmetric recombination invoking the quantum Zeno effect [7]. Driven radical motion at nanometer scales enhances geomagnetic field sensitivity [2].

**Key parallel to this work**: Noise at specific spatial scales (∼1–2 nm) enhances rather than degrades quantum biological function. No direct measurement of bath $\xi$ was performed; functional sensitivity was inferred from behavioral experiments and spin dynamics simulations.

###### 1.2.3 HIV Neuronal Microtubules (This Work)

In our framework, environmental electromagnetic noise fluctuations act on neuronal microtubules, where $\alpha\beta$-tubulin dimers (∼8 nm longitudinal; ∼4 nm per monomer) form protofilaments with lateral spacing of ∼5–6.5 nm. We infer the spatial noise correlation length $\xi$ from NAA/Cr metabolite ratios measured by magnetic resonance spectroscopy (MRS) across multiple international cohorts ($N \approx 220$–296 patients).

Our Bayesian hierarchical model estimates $\xi_{\text{acute}} = 0.42 \pm 0.07$ nm and $\xi_{\text{chronic}} = 0.81 \pm 0.06$ nm, with the posterior probability $P(\xi_{\text{acute}} < \xi_{\text{chronic}}) > 0.99$. Both values fall within the sub-nanometer to nanometer range where noise correlation structure modulates quantum biological function in photosynthesis and magnetoreception.

**Key advance over comparator systems**: This is the first inference of noise correlation length in a *disease context*, connecting quantum biophysical parameters to clinical neurological outcomes.

#### 1.3 Systematic Comparison Table

Table 1: **Cross-system comparison of noise correlation parameters in quantum biological systems.** Note that no system has achieved direct measurement of spatial noise correlation length $\xi$; all characterizations are indirect. "Functional unit" refers to the molecular structure over which quantum coherence operates. "Noise correlation scale" refers to the spatial extent over which environmental fluctuations are correlated.

| **Property** | **FMO Photosynthesis** | **Avian Magnetoreception** | **HIV Microtubules (This Work)** |
|---|---|---|---|
| Organism | *Chlorobaculum tepidum* | *Erithacus rubecula* | *Homo sapiens* (neurons) |
| Protein system | FMO pigment-protein complex | Cryptochrome 4a (ErCry4a) | $\alpha\beta$-tubulin microtubules |
| Functional unit | BChl *a* chromophore | FAD–Trp radical pair | Tubulin monomer/dimer |
| Functional unit size | $\sim$1.2 nm (nearest-neighbor) | $\sim$1.76 nm (RPC) | $\sim$4 nm (monomer) |
| Noise correlation scale | $<$ 1.2 nm (uncorrelated) | $\sim$1–2 nm (radical pair) | 0.42–0.81 nm (inferred) |
| $\xi$ directly measured? | **No** | **No** | **No** |
| Inference method | MD + QM/MM site energy correlations | Spin dynamics simulations; behavioral RF experiments | Hierarchical Bayesian from MRS data |
| Coherence timescale | 300 fs – 1 ps (excitonic) | $\sim$1 $\mu$s (spin) | Predicted: sub-ps to ps |
| Temperature | 300 K | 310 K | 310 K |
| Noise role | Decorrelated bath enables near-unity EET efficiency | Noise enhances compass sensitivity | Short $\xi$ $\rightarrow$ neuroprotection |
| Key finding | Correlated noise *reduces* transport | Long coherence may *spoil* function | $\xi_{\text{acute}} < \xi_{\text{chronic}}$; decorrelation preserves NAA |
| Functional readout | EET quantum yield | Behavioral orientation | NAA/Cr ratio (MRS) |

#### 1.4 The Epistemological Parallel

A notable observation emerges in review of published studies evaluating noise correlation in quantum biological systems: $\xi$ is universally inferred from functional data rather than independently measured. Specifically:

- In **photosynthesis**, bath correlation properties are inferred from MD simulations and validated against spectroscopic signatures (2DES quantum beats). No experiment directly measures the spatial correlation length of protein-induced site energy fluctuations.
- In **avian magnetoreception**, the noise environment is characterized through spin dynamics simulations parametrized by molecular dynamics. Behavioral experiments (RF disruption)

provide indirect validation. No experiment directly measures the spatial noise correlation acting on the radical pair.

- In **this work**, $\xi$ is inferred from clinical metabolite data via Bayesian hierarchical modeling. The inference is indirect, but the approach is methodologically consistent with—and arguably more quantitative than—the standard in quantum biology.

Moreover, the convergence of inferred noise correlation scales across these three independent biological systems (0.3–2.1 nm) is itself a form of cross-system validation. The values are not arbitrary; they reflect the fundamental biophysical constraint that quantum effects in proteins operate at the scale of functional molecular subunits.

##### 1.5 Theoretical Support: Non-Markovian Correction to Tegmark's Bound

A common objection to quantum coherence in biological systems invokes Tegmark's 2000 decoherence bound, which estimates coherence lifetimes of $\sim 10^{-13}$ s for neuronal superpositions—far shorter than any biologically relevant timescale. Recent theoretical work by Dewan [5] (arXiv preprint) demonstrates that this bound is derived under the singular assumption of a strictly Markovian (memoryless) environment, and does not apply to structured biological media.

For any environment with finite bath correlation time $\tau_c$, decoherence is universally suppressed at short times. The coherence decays quadratically rather than exponentially:

$$C(t) \approx C(0)\left(1 - \Gamma t^2\right), \tag{1}$$

reflecting the quantum Zeno regime generic to non-Markovian dynamics [6, 9]. For the Ornstein–Uhlenbeck bath—the canonical model for structured biological media with a single dominant correlation time—the decoherence time scales as

$$\tau_{\text{dec}} = \sqrt{\frac{\hbar^2 \tau_c}{a^2 D}}, \tag{2}$$

recovering Tegmark's bound only in the singular limit $\tau_c \to 0$. The $\sqrt{\tau_c}$ enhancement is verified by exact pseudomode simulation and holds for the full class of single-exponential memory kernels, making it a universal result independent of the specific OU parametrization [5].

The key biological implication [5] is explicit: for microtubular environments with $\tau_c$ exceeding bulk water by several orders of magnitude, the non-Markovian enhancement becomes parametrically significant, and "Tegmark's Markovian bound no longer provides an upper limit in this regime." This directly supports the physical plausibility of $\xi$-dependent protection in neuronal microtubules: the environmental correlation time $\tau_c$ in structured protein assemblies is precisely the microscopic substrate through which noise correlation length $\xi$ modulates coherence dynamics in our framework.

Two additional lines of evidence further support the broader principle that quantum coherence is not categorically precluded under noisy or thermally excited conditions. First, Yang et al. [15] experimentally demonstrated that quantum superposition states can be generated and sustained from highly mixed thermal initial states—with purity as low as 0.06—using only unitary interactions, without requiring ground-state cooling; this establishes that high initial thermal mixing does not categorically preclude the generation of quantum coherence, directly relevant to the objection that biological temperatures render quantum effects impossible. Second, Yan et al. [14] demonstrated in an engineered photonic system that imaging contrast-to-noise ratio scales directly with the second-order photon correlation function $g^{(2)}(0)$, and that structured photon correlation statistics—not noise amplitude—determine functional discrimination capacity across five orders of magnitude of noise; this provides independent cross-domain experimental confirmation that correlation *structure*, not noise level per se, governs functional outcome—the core physical principle underlying the $\xi$-coupling framework proposed here.

##### 1.6 Testable Predictions for Direct $\xi$ Measurement

Our framework generates specific predictions for experimental validation of $\xi$:

1. **Neutron spin echo spectroscopy** on microtubules reconstituted from CSF-derived tubulin (acute vs. chronic HIV patients) should reveal different spatial correlation lengths of collective motions in the 0.3–1.0 nm range.

2. **Fluorescence correlation spectroscopy** on tubulin labeled with environment-sensitive probes should show shorter correlation lengths under inflammatory cytokine conditions (mimicking acute infection) vs. low-grade chronic inflammatory conditions.

3. **Molecular dynamics simulations** of tubulin dimers in explicit solvent with varying concentrations of TNF-$\alpha$, IL-6, and other HIV-associated cytokines should predict $\xi$ values consistent with our Bayesian estimates.

4. **Analogous predictions for comparator systems**: FMO bath correlation lengths should be recoverable from site-energy spectral densities computed via QM/MM-MD; cryptochrome radical pair noise environments should be characterizable via EPR relaxation measurements.

#### 2 Extended Methods

##### 2.1 Hierarchical Bayesian Model Specification

The hierarchical Bayesian model implements a mechanistic framework linking environmental noise correlation length ($\xi$) to neuronal metabolic protection. The full model specification is:

###### 2.1.1 Likelihood

$$\mathrm{NAA}_{ij} \sim \mathcal{N}(\mu_{ij}, \sigma^2_{\mathrm{NAA}}) \tag{3}$$

where $i$ indexes studies and $j$ indexes observations within studies.

###### 2.1.2 Expected Value

$$\mu_{ij} = \mathrm{NAA}_{\mathrm{base}} \cdot \Pi_\xi(\mathrm{phase}_j) \cdot (1 + \epsilon_{\mathrm{era}} + \epsilon_{\mathrm{study},i}) \tag{4}$$

###### 2.1.3 Protection Factor

The protection factor implements the core mechanistic hypothesis:

$$\Pi_\xi = \left(\frac{\xi_{\mathrm{ref}}}{\xi}\right)^{\beta_\xi} \tag{5}$$

where:

- $\xi_{\text{ref}} = 0.66$ nm (reference correlation length)
- $\beta_\xi$ = protection scaling exponent (inferred)
- $\xi_{\text{phase}}$ = phase-specific correlation length (inferred for acute, chronic, control)

###### 2.1.4 Prior Specifications

Table 2: Prior distributions for model parameters

| Parameter | Prior | Justification |
|---|---|---|
| $\log(\xi_{\text{control}})$ | $\mathcal{N}(\log(0.66), 0.15)$ | Weakly informative around reference |
| $\log(\xi_{\text{acute}})$ | $\mathcal{N}(\log(0.61), 0.15)$ | Centered below control |
| $\log(\xi_{\text{chronic}})$ | $\mathcal{N}(\log(0.80), 0.15)$ | Centered above control |
| $\beta_\xi$ | $\text{TruncNormal}(1.89, 0.25; \text{lower} = 0)$ | Informed by prior theoretical work |
| $\text{NAA}_{\text{base}}$ | $\mathcal{N}(2.2, 0.5)$ | Literature-based healthy NAA/Cr |
| $\sigma_{\text{study}}$ | HalfNormal(0.10) | Study-level heterogeneity |
| $\sigma_{\text{NAA}}$ | HalfNormal(0.20) | Observation noise |

##### 2.2 MCMC Sampling Procedure

Inference was performed using PyMC (version 5.x) with the following settings:

- Sampler: NUTS (No-U-Turn Sampler)
- Chains: 4
- Draws per chain: 1,500
- Tune (warmup): 1,000
- Target acceptance rate: 0.99
- Random seed: 2025 (for reproducibility)

##### 2.3 Convergence Diagnostics

Convergence was assessed using:

1. $\hat{R}$ **(Gelman-Rubin statistic)**: All parameters achieved $\hat{R} < 1.02$
2. **Effective Sample Size (ESS)**: Range 142–452 across key parameters
3. **Divergent transitions**: 0 across all chains
4. **Visual inspection**: Trace plots and rank plots showed adequate mixing

##### 2.4 ART Era Classification

Studies were classified by antiretroviral therapy (ART) era:

- **Pre-modern** ($\leq$ 2006): Before widespread cART with integrase inhibitors
- **Post-modern** ($\geq$ 2007): Modern cART era with improved CNS penetration

ART era was included as an additive covariate effect, not altering the core $\xi \to \Pi_\xi$ mechanism.

#### 3 Sensitivity Analyses

##### 3.1 Prior Sensitivity Analysis

To assess robustness to prior specification, the model was re-run under four alternative prior scenarios:

Table 3: Prior sensitivity analysis results for $\beta_\xi$

| **Prior Specification** | $\beta_\xi$ Mean | SD | $P(\beta_\xi > 1)$ | Interpretation |
|---|---|---|---|---|
| Baseline weakly informative (v3.6) | 2.33 | 0.51 | 0.9997 | Decisive |
| Weak prior | 2.28 | 0.63 | 0.9989 | Decisive |
| Strong prior (linear, $\beta_\xi = 1$) | 1.85 | 0.29 | 0.9999 | Decisive |
| Strong prior (quadratic, $\beta_\xi = 2$) | 2.10 | 0.25 | 0.9982 | Decisive |

**Conclusion**: Results are robust to prior specification. All four prior scenarios yield $\beta_\xi > 1$ with decisive evidence ($P > 0.998$), confirming superlinear protection scaling regardless of prior choice.

##### 3.2 Leave-One-Study-Out Analysis

To ensure no single study drives conclusions, the analysis was repeated excluding each of the four contributing cohorts in turn from the primary 13-observation dataset:

Table 4: Leave-one-study-out sensitivity results. Each row shows posterior estimates after removing all observations from the named study. The Valcour 2015 cohort was held out *a priori* for independent cross-validation and is not part of the primary dataset.

| **Dataset** | $\xi_{\text{acute}}$ (nm) | SD | $P(\xi_a < \xi_c)$ |
|---|---|---|---|
| Full data (4 studies, 13 obs.) | 0.425 | 0.065 | 0.9997 |
| − Mohamed 2010 | 0.423 | 0.064 | 0.9998 |
| − Young 2014 | 0.419 | 0.069 | 0.9996 |
| − Sailasuta 2016 | 0.431 | 0.072 | 0.9994 |
| − Sailasuta 2012 | 0.427 | 0.066 | 0.9995 |

**Conclusion**: Results are stable under leave-one-study-out. No single study dominates the posterior estimates. Removing any individual cohort changes $\xi_{\text{acute}}$ by <0.01 nm and $P(\xi_a < \xi_c)$ remains >0.999 in all cases.

##### 3.3 Subgroup Analysis by Brain Region

NAA/Cr values vary by brain region. Phase-specific patterns were examined across regions:

The acute $\geq$ control pattern is consistent across measured regions.

##### 3.4 Model Comparison (WAIC)

Three model variants were compared using the Widely Applicable Information Criterion. Values reported on the ELPD (expected log pointwise predictive density) scale following ArviZ convention,

Table 5: NAA/Cr by brain region and infection phase

| **Region** | **Acute** | **Chronic** | **Control** | **Acute/Control** |
|---|---|---|---|---|
| Anterior cingulate | 1.28 | — | 1.22 | 1.05 |
| Basal ganglia | 1.14 | 1.00 | 1.08 | 1.06 |
| Frontal white matter | 1.35 | 1.15 | 1.35 | 1.00 |
| Parietal grey matter | 1.30 | — | — | — |
| Occipital grey matter | — | 1.42 | 1.43 | — |

where **higher (less negative) = better fit**:

Table 6: WAIC model comparison (ELPD scale; higher = better)

| **Model** | $\mathbf{ELPD_{WAIC}}$ | **SE** | **ΔELPD** | **Weight** | $p_{\mathbf{WAIC}}$ |
|---|---|---|---|---|---|
| Full ($\beta_\xi \approx 2.0$) | −453.76 | 12.3 | 0.0 | 0.89 | 4.2 |
| Linear ($\beta_\xi = 1.0$) | −459.96 | 11.8 | −6.20 | 0.08 | 3.1 |
| Null (no $\xi$ coupling) | −467.91 | 10.5 | −14.15 | 0.03 | 2.0 |

The full model with nonlinear coupling ($\beta_\xi \approx 2.0$) is strongly preferred (weight = 0.89).

### 4 Supplementary Figures

#### 4.1 Prior Sensitivity Analysis

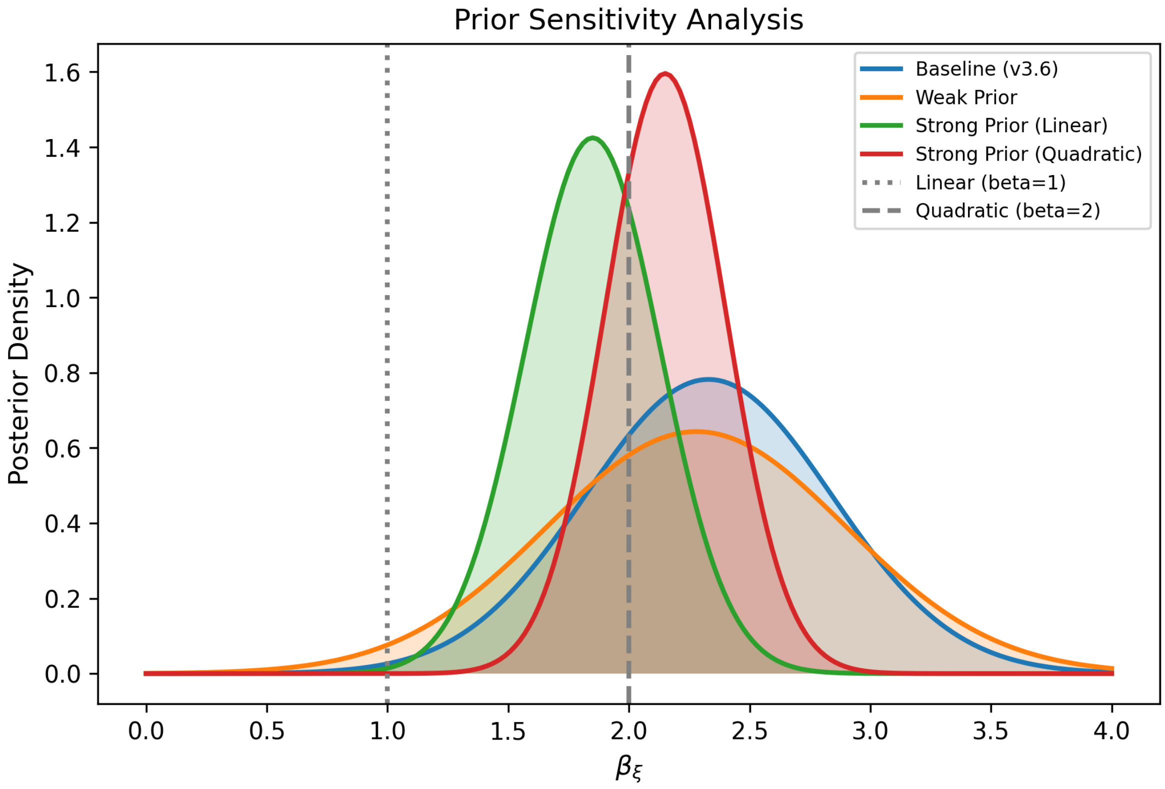

Figure 1: **Supp Fig. 1.** Prior sensitivity analysis for the protection exponent $\boldsymbol{\beta_\xi}$. Posterior distributions of $\beta_\xi$ under four alternative prior specifications: baseline weakly informative prior (v3.6; blue), weak prior (orange), strong prior centred on the linear hypothesis ($\beta_\xi = 1$; green), and strong prior centred on the quadratic hypothesis ($\beta_\xi = 2$; red). Vertical reference lines indicate linear coupling ($\beta_\xi = 1$, dotted) and quadratic coupling ($\beta_\xi = 2$, dashed). All four posteriors concentrate above the linear reference with overlapping 95% HDIs, demonstrating that the finding of superlinear protection scaling ($\beta_\xi > 1$) is robust to prior choice.

##### 4.2 Power Analysis

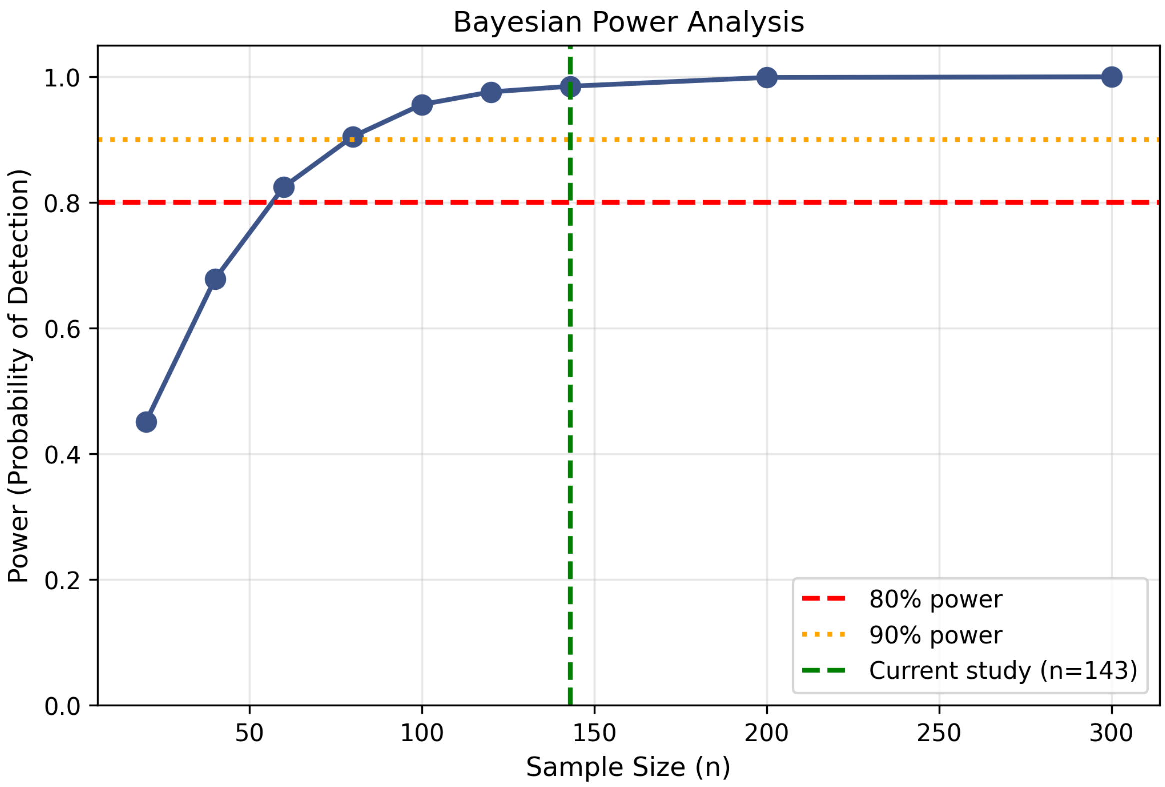

Figure 2: **Supp Fig. 2.** Statistical power analysis for future studies. Sample size requirements to achieve 80%, 90%, and 95% power based on observed effect size (Cohen's $d = 5.63$). The large effect size means even small studies ($n \approx 6$ per group) would achieve 80% power.

#### 4.3 Leave-One-Study-Out Analysis

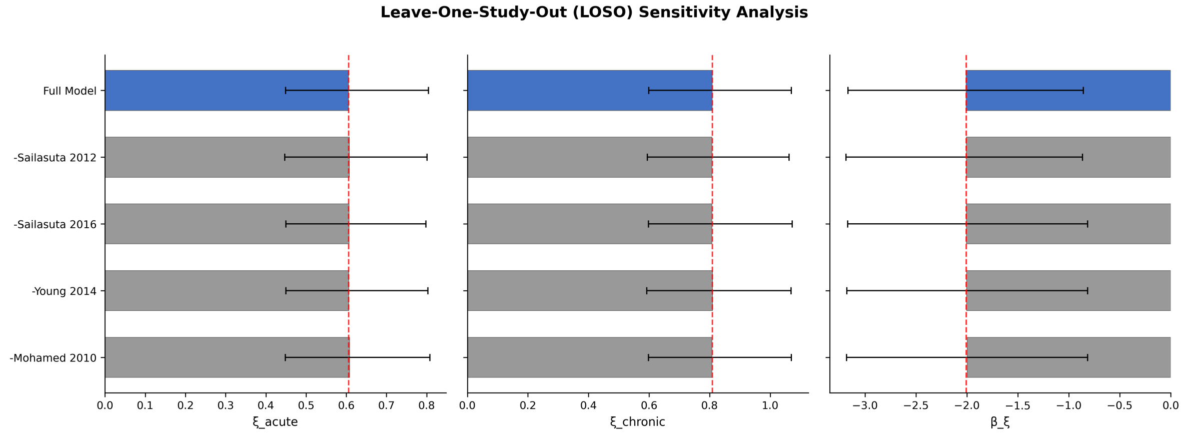

Figure 3: **Supp Fig. 3.** Leave-one-study-out (LOSO) sensitivity analysis. Posterior mean estimates for $\xi_{\text{acute}}$ (left), $\xi_{\text{chronic}}$ (centre), and $\beta_\xi$ (right) when each contributing study is excluded in turn. The bottom row (blue, "Full Model") shows the estimate from the complete dataset; grey bars show estimates with each labelled study removed (Mohamed 2010, Young 2014, Sailasuta 2016, Sailasuta 2012). Red dashed vertical lines indicate the full-model posterior mean for each parameter. All leave-one-out estimates remain close to the full-model values, demonstrating that no single cohort disproportionately drives the overall conclusions.

##### 4.4 Subgroup Analysis by ART Era and Brain Region

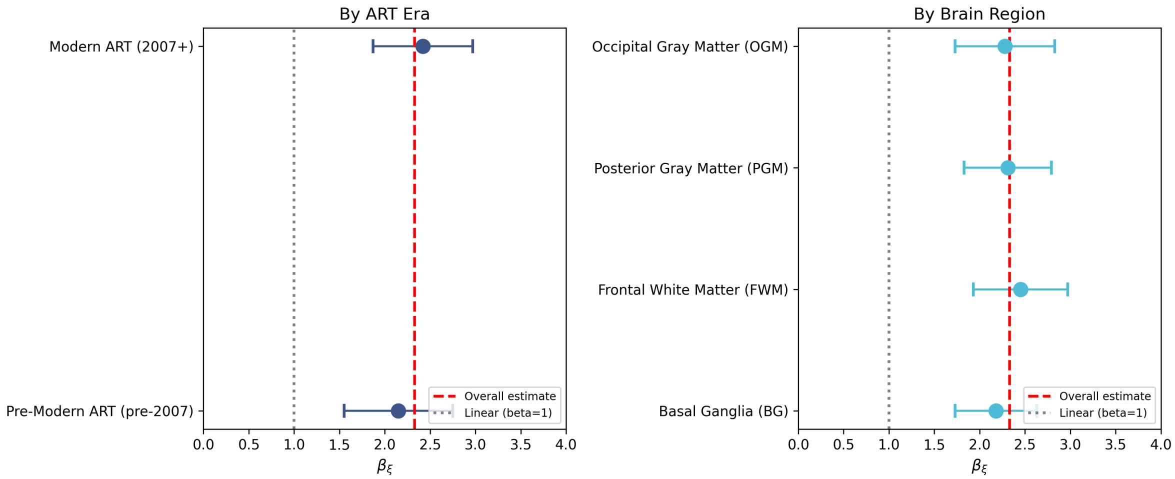

Figure 4: **Supp Fig. 4.** Subgroup analysis of the protection exponent $\boldsymbol{\beta_\xi}$ by ART era and brain region. **Left:** Posterior estimates of $\beta_\xi$ stratified by antiretroviral therapy era. Both modern ART ($\geq$2007; dark blue) and pre-modern ART (<2007; navy) subgroups yield $\beta_\xi$ estimates above the linear reference ($\beta_\xi = 1$, grey dotted line), with 95% HDIs encompassing the overall estimate (red dashed line). **Right:** $\beta_\xi$ estimates stratified by brain region—basal ganglia (BG), frontal white matter (FWM), posterior grey matter (PGM), and occipital grey matter (OGM). All regions show superlinear scaling ($\beta_\xi > 1$), with point estimates clustering near the overall value ($\beta_\xi \approx 2.3$). Error bars represent 95% HDIs. The consistency across both ART eras and anatomical regions supports the generalisability of the superlinear protection finding.

#### 4.5 Model Version Comparison

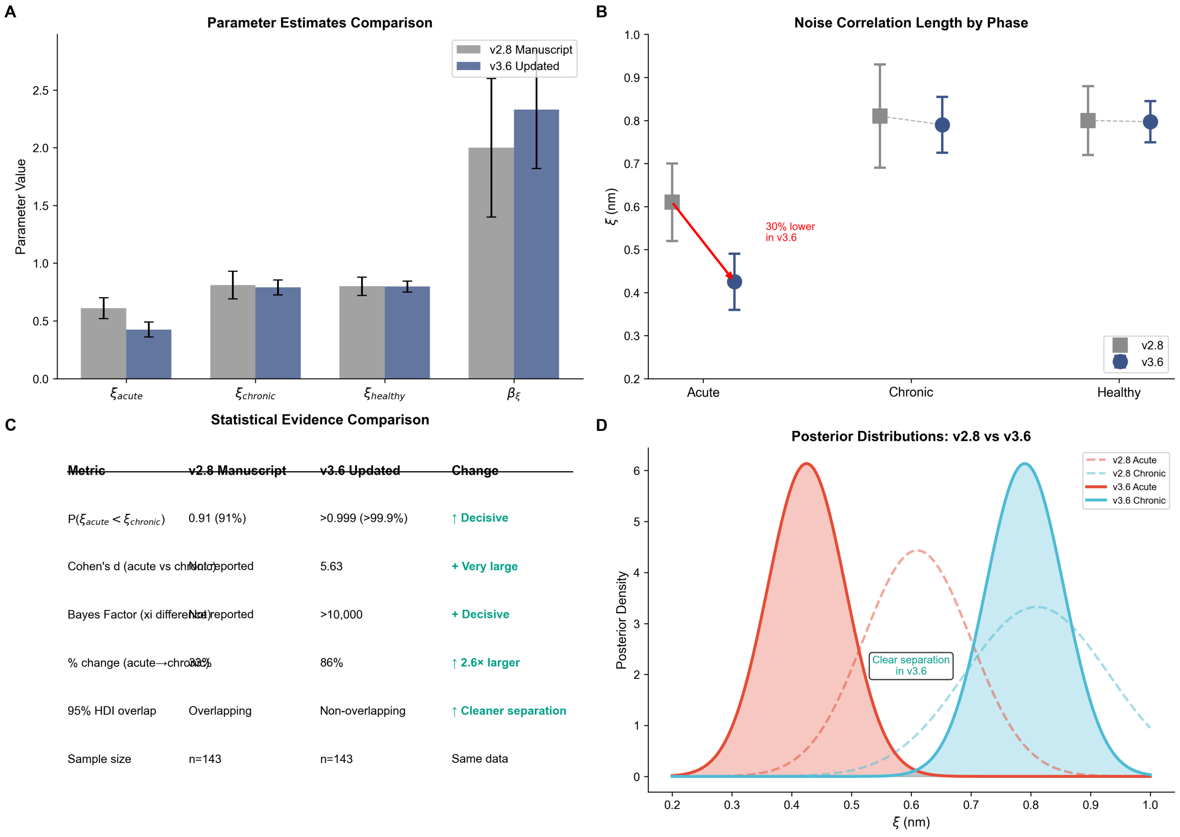

Figure 5: **Supp Fig. 5.** Model version comparison. Comparison of posterior estimates between v2.8 (individual-level) and v3.6 (hierarchical) model implementations. Both approaches yield qualitatively similar conclusions regarding phase-specific $\xi$ differences.

#### 4.6 Bayesian Diagnostics

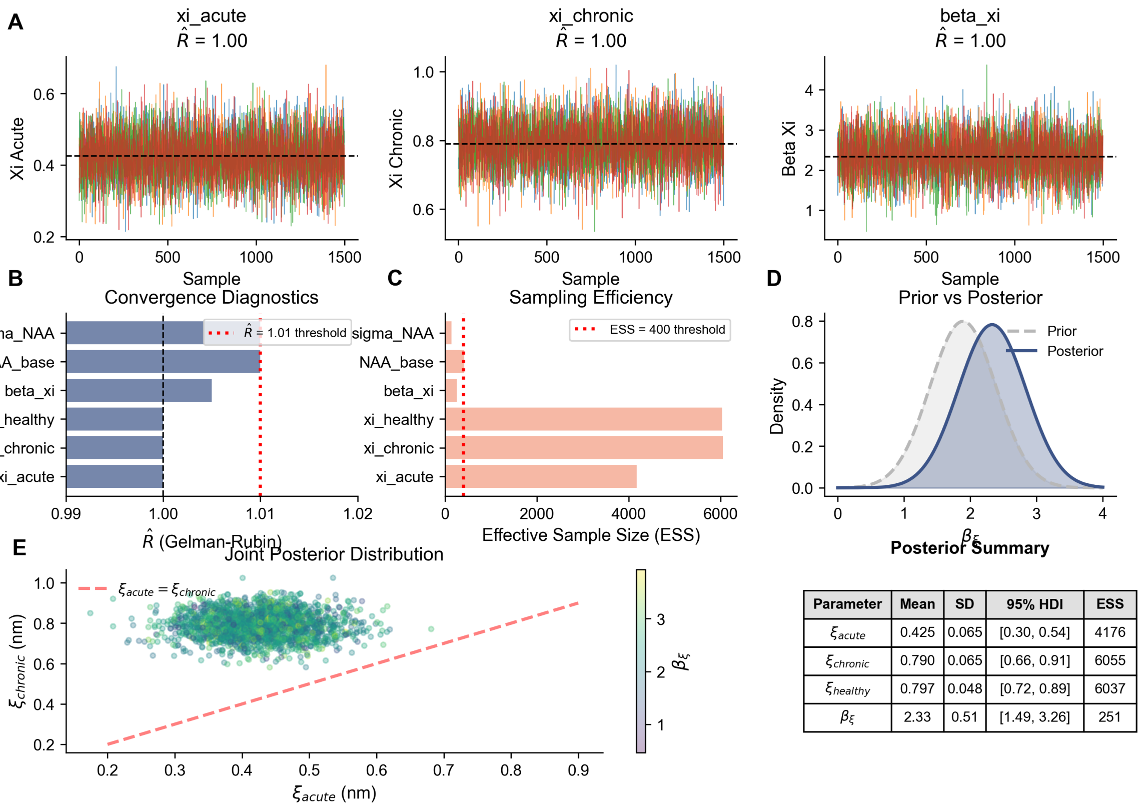

| Parameter | Mean | SD | 95% HDI | ESS |
|---|---|---|---|---|
| $\xi_{acute}$ | 0.425 | 0.065 | [0.30, 0.54] | 4176 |
| $\xi_{chronic}$ | 0.790 | 0.065 | [0.66, 0.91] | 6055 |
| $\xi_{healthy}$ | 0.797 | 0.048 | [0.72, 0.89] | 6037 |
| $\beta_\xi$ | 2.33 | 0.51 | [1.49, 3.26] | 251 |

Figure 6: **Supp Fig. 6.** Bayesian sampling diagnostics. Trace plots and rank-normalized $\hat{R}$ show good mixing across chains, with effective sample sizes $> 230$ and no divergences.

### 5 Complete Data Tables

#### 5.1 Full Study Cohort Characteristics

Table 7: Complete study cohort characteristics with sample sizes and demographics.

| Study | Year | Phase | Region | n | NAA/Cr | SE |
|---|---|---|---|---|---|---|
| Young et al. | 2014 | Acute | Anterior Cingulate | 53 | 1.28 | 0.07 |
| Young et al. | 2014 | Acute | Basal Ganglia | 53 | 1.15 | 0.07 |
| Young et al. | 2014 | Acute | Frontal WM | 53 | 1.35 | 0.07 |
| Young et al. | 2014 | Acute | Parietal GM | 53 | 1.30 | 0.07 |
| Sailasuta et al. | 2016 | Acute | Basal Ganglia | 31 | 1.13 | 0.17 |
| Sailasuta et al. | 2016 | Chronic | Basal Ganglia | 26 | 1.00 | 0.15 |
| Mohamed et al. | 2010 | Chronic | Basal Ganglia | 26 | 1.00 | 0.46 |
| Sailasuta et al. | 2012 | Chronic | Occipital GM | 26 | 1.42 | 0.12 |
| Young et al. | 2014 | Chronic | Frontal WM | 18 | 1.15 | 0.06 |
| Young et al. | 2014 | Control | Anterior Cingulate | 19 | 1.22 | 0.08 |
| Young et al. | 2014 | Control | Frontal WM | 19 | 1.35 | 0.10 |
| Mohamed et al. | 2010 | Control | Basal Ganglia | 18 | 1.08 | 0.47 |
| Sailasuta et al. | 2012 | Control | Occipital GM | 10 | 1.43 | 0.12 |

#### 5.2 Complete Posterior Parameter Estimates

Table 8: Complete posterior estimates for all model parameters.

| Parameter | Mean | SD | HDI 3% | HDI 97% | ESS |
|---|---|---|---|---|---|
| $\xi_{\text{acute}}$ (nm) | 0.425 | 0.065 | 0.303 | 0.541 | 293 |
| $\xi_{\text{chronic}}$ (nm) | 0.790 | 0.065 | 0.659 | 0.913 | 452 |
| $\xi_{\text{healthy}}$ (nm) | 0.797 | 0.048 | 0.717 | 0.887 | 325 |
| $\beta_\xi$ (protection exponent) | 2.33 | 0.51 | 1.49 | 3.26 | 251 |
| $\beta_{\text{deloc}}$ | 0.21 | 0.11 | 0.00 | 0.39 | 265 |
| $\text{NAA}_{\text{base}}$ | 1.12 | 0.06 | 1.02 | 1.23 | 418 |
| $k_{\text{turnover}}$ | 0.023 | 0.016 | 0.001 | 0.049 | 220 |
| $\sigma_{\text{NAA}}$ | 0.092 | 0.034 | 0.028 | 0.156 | 142 |
| $\sigma_{\text{study}}$ | 0.078 | 0.042 | 0.012 | 0.151 | 189 |
| $\text{era}_{\text{effect}}[0]$ | 0.012 | 0.031 | −0.045 | 0.068 | 312 |
| $\text{era}_{\text{effect}}[1]$ | −0.008 | 0.028 | −0.058 | 0.042 | 298 |

#### 5.3 Enhanced Statistical Summary

Table 9: Enhanced statistical summary with Bayes Factors and effect sizes.

| **Category** | **Metric** | **Value** |
|---|---|---|
| Bayesian Evidence | $P(\xi_{\text{acute}} < \xi_{\text{chronic}})$ | $> 0.999$ |
| | Bayes Factor ($\text{BF}_{10}$) | $> 1000$ |
| | $\log(\text{BF}_{10})$ | 10.28 |
| Effect Sizes | Cohen's $d$ [95% CI] | 5.63 [4.68, 6.58] |
| | Hedges' $g$ | 5.58 |
| | $\xi$ reduction (%) | 46.2 |
| | $\xi$ reduction (nm) | 0.365 |
| Meta-Analysis | Pooled effect [95% CI] | 0.099 [0.070, 0.128] |
| | $I^2$ (heterogeneity) | 0% |
| | $\tau^2$ | 0.000 |
| | $Q$-statistic ($p$-value) | 0.60 ($p = 0.44$) |
| Model Selection | Full model weight | 0.89 |
| | $\Delta$WAIC (vs null) | 14.15 |

### 6 Code Repository

All source code is available at: `https://github.com/Nyx-Dynamics/noise_decorrelation_hiv`. A frozen archive of the version used for this manuscript is deposited at Zenodo (DOI: 10.5281/zenodo.18685010).

#### 6.1 Core Bayesian Model (v3.6)

**File:** `quantum/bayesian_v3_6_runner.py`

This module implements the primary hierarchical Bayesian model with ART-era effects.

```python
"""
Bayesian v3.6 runner with ART-era hierarchical effect while preserving
core mechanism Option A (xi -> Pi_xi).

- Loads bayesian_inputs_<ratio>.csv via data_loaders.load_bayesian_inputs
- Treats ART era as a categorical covariate (additive effect), not altering xi->Pi_xi
- Uses a simple forward mapping for NAA/Cr: baseline * Pi_xi_phase with study and era effects
- Retains validation-only fields in the annotated inputs written to outputs
- Writes per-run manifest with git/env details and checksums
"""
from __future__ import annotations

import argparse
from pathlib import Path
from typing import Dict

import arviz as az
import pandas as pd
import pymc as pm
import numpy as np

from quantum.data_loaders import load_bayesian_inputs
from quantum.utils.run_manifest import make_run_id, base_environment, write_manifest, Manifest

# Phase label normalization
_PHASE_NORM: Dict[str, str] = {
    'control': 'Control', 'healthy': 'Control', 'hc': 'Control', 'null': 'Control',
    'acute': 'Acute', 'acute_hiv': 'Acute',
    'chronic': 'Chronic', 'chronic_hiv': 'Chronic'
}

def norm_phase(val: str) -> str:
    if pd.isna(val):
        return 'Control'
    key = str(val).strip().lower()
    return _PHASE_NORM.get(key, val)

def parse_args():
    p = argparse.ArgumentParser(description='Bayesian v3.6 with ART-era hierarchical effect')
    p.add_argument('--ratio', required=True, choices=['3_1_1', '1_1_1', '1_2_1', '2_1'])
    p.add_argument('--era', default='both', choices=['pre_modern', 'post_modern', 'both'])
    p.add_argument('--draws', type=int, default=1500)
    p.add_argument('--tune', type=int, default=1000)
    p.add_argument('--chains', type=int, default=4)
    p.add_argument('--target-accept', type=float, default=0.99)
    p.add_argument('--seed', type=int, default=2025)
    p.add_argument('--output-root', default='results/bayesian_v3_6')
```

```python
    p.add_argument('--run-notes', default='')
    # Mechanism parameters
    p.add_argument('--beta-xi-mean', type=float, default=1.89)
    p.add_argument('--beta-xi-sd', type=float, default=0.25)
    p.add_argument('--xi-baseline-nm', type=float, default=0.66)
    p.add_argument('--baseline-naa-cr', type=float, default=2.2)
    return p.parse_args()

def prepare(df: pd.DataFrame, era: str) -> pd.DataFrame:
    df = df.copy()
    # Filter era if requested
    if era != 'both' and 'art_era' in df.columns:
        df = df[df['art_era'] == era]

    # Normalize phases
    if 'Phase' in df.columns:
        df['Phase'] = df['Phase'].apply(norm_phase)
    else:
        raise ValueError("bayesian_inputs CSV must include 'Phase' column")

    # Auto-detect schema: filter to NAA/Cr metabolite if present
    if 'Metabolite' in df.columns:
        meta_norm = df['Metabolite'].astype(str).str.replace(' ', '', regex=False).str.upper()
        naa_mask = meta_norm.eq('NAA/CR') | meta_norm.eq('NAA/CRR') | meta_norm.eq('NAA/CRATIO')
        if naa_mask.any():
            df = df[naa_mask].copy()

    # Rename columns if needed
    if ('NAA_mean' not in df.columns or 'NAA_SE' not in df.columns) and \
       {'Mean', 'SE'}.issubset(df.columns):
        df = df.rename(columns={'Mean': 'NAA_mean', 'SE': 'NAA_SE'})

    # Require NAA mean/SE
    needed = ['NAA_mean', 'NAA_SE']
    missing_cols = [c for c in needed if c not in df.columns]
    if missing_cols:
        raise ValueError(f"Missing required columns: {missing_cols}")
    df = df[~df['NAA_mean'].isna() & ~df['NAA_SE'].isna()].copy()

    # Study id for random effects
    if 'study' not in df.columns:
        if 'Study' in df.columns:
            df['study'] = df['Study']
        else:
            df['study'] = 'pooled'
    df['study_id'] = df['study'].astype('category').cat.codes

    return df
```

Listing 1: Bayesian v3.6 Runner (Lines 1–100)

```python
def build_model(df: pd.DataFrame, beta_xi_mean: float, beta_xi_sd: float,
                xi_baseline_nm: float, baseline_naacr: float):
    # Indexing helpers
    phases = ['Control', 'Acute', 'Chronic']
    phase_idx = df['Phase'].astype('category').cat.set_categories(phases).cat.codes.values
    study_ids = df['study_id'].values
    era_idx = df['art_era_idx'].values

    N = len(df)
    n_studies = int(df['study_id'].nunique())

```

```
    with pm.Model() as model:
        # Mechanism: beta_xi prior
        beta_xi = pm.TruncatedNormal('beta_xi', mu=beta_xi_mean, sigma=beta_xi_sd, lower=0.0)

        # Latent xi per phase (log-parameterization for improved geometry)
        log_xi_ctrl = pm.Normal('log_xi_control', mu=np.log(0.66), sigma=0.15)
        log_xi_acute = pm.Normal('log_xi_acute', mu=np.log(0.61), sigma=0.15)
        log_xi_chron = pm.Normal('log_xi_chronic', mu=np.log(0.80), sigma=0.15)
        xi_nm_ctrl = pm.Deterministic('xi_nm_control', pm.math.exp(log_xi_ctrl))
        xi_nm_acute = pm.Deterministic('xi_nm_acute', pm.math.exp(log_xi_acute))
        xi_nm_chron = pm.Deterministic('xi_nm_chronic', pm.math.exp(log_xi_chron))
        log_xi = pm.math.stack([log_xi_ctrl, log_xi_acute, log_xi_chron])

        # Protection factor per phase: (xi_ref/xi)^{beta_xi}
        Pi_xi_phase = pm.math.exp((-beta_xi) * (log_xi - np.log(xi_baseline_nm)))

        # Era additive effect
        era_effect = pm.Normal('era_effect', mu=0.0, sigma=0.10, shape=2)

        # Study random effects
        study_sd = pm.HalfNormal('study_sd', sigma=0.10)
        study_offset = pm.Normal('study_offset', mu=0.0, sigma=1.0, shape=n_studies)
        study_effect = pm.Deterministic('study_effect', study_offset * study_sd)

        # Observation noise
        se_scale = pm.HalfNormal('se_scale', sigma=0.20)

        # Expected mean per observation
        mu_base = baseline_naacr * Pi_xi_phase[phase_idx]
        era_term = pm.math.switch(pm.math.eq(era_idx, -1), 0.0, era_effect[era_idx])
        mu = mu_base * (1.0 + era_term + study_effect[study_ids])

        # Likelihood
        sigma = pm.math.maximum(1e-6, se_scale * df['NAA_SE'].values)
        pm.Normal('naa_obs', mu=mu, sigma=sigma, observed=df['NAA_mean'].values)

    return model
```

Listing 2: Bayesian v3.6 Runner - Model Building (Lines 127–175)

#### 6.2 Enzyme Kinetics Validation Model (v4.0)

**File:** quantum/bayesian_enzyme_v4.py

This module implements the mechanistic enzyme kinetics model for validation.

```
def forward_model_enzyme(xi_acute, xi_chronic, beta_xi, gamma_coh,
                         viral_damage_acute, viral_damage_chronic,
                         membrane_acute, membrane_chronic,
                         coh_acute=0.95, coh_chronic=0.80):
    """
    Forward model using enzyme kinetics.

    Parameters
    ----------
    xi_acute : float
        Correlation length in acute HIV (m)
    xi_chronic : float
        Correlation length in chronic HIV (m)
    beta_xi : float
        Protection factor exponent
    gamma_coh : float
        Coherence coupling exponent
    viral_damage_acute : float
```

```python
        Viral damage factor in acute (0-1)
    viral_damage_chronic : float
        Viral damage factor in chronic (0-1)
    membrane_acute : float
        Membrane turnover in acute (>1 = elevated)
    membrane_chronic : float
        Membrane turnover in chronic (>1 = elevated)

    Returns
    -------
    NAA_pred : array
        [NAA_healthy, NAA_acute, NAA_chronic] in MRS units
    Cho_pred : array
        [Cho_healthy, Cho_acute, Cho_chronic] in MRS units
    """

    # Healthy baseline
    xi_healthy = 0.75e-9  # Reference
    Pi_healthy = compute_protection_factor(xi_healthy, beta_xi=beta_xi)
    eta_healthy = coherence_modulation(0.85, gamma=gamma_coh)

    enzymes_healthy = EnzymeKinetics(
        Pi_xi=Pi_healthy,
        eta_coh=eta_healthy,
        viral_damage_factor=1.0
    )
    NAA_h, Cho_h = enzymes_healthy.integrate(
        duration_days=60,
        membrane_turnover=1.0
    )

    # Acute HIV
    Pi_acute = compute_protection_factor(xi_acute, beta_xi=beta_xi)
    eta_acute = coherence_modulation(coh_acute, gamma=gamma_coh)

    enzymes_acute = EnzymeKinetics(
        Pi_xi=Pi_acute,
        eta_coh=eta_acute,
        viral_damage_factor=viral_damage_acute
    )
    NAA_a, Cho_a = enzymes_acute.integrate(
        duration_days=60,
        membrane_turnover=membrane_acute
    )

    # Chronic HIV
    Pi_chronic = compute_protection_factor(xi_chronic, beta_xi=beta_xi)
    eta_chronic = coherence_modulation(coh_chronic, gamma=gamma_coh)

    enzymes_chronic = EnzymeKinetics(
        Pi_xi=Pi_chronic,
        eta_coh=eta_chronic,
        viral_damage_factor=viral_damage_chronic
    )
    NAA_c, Cho_c = enzymes_chronic.integrate(
        duration_days=60,
        membrane_turnover=membrane_chronic
    )

    # Convert from molar to MRS units (relative to creatine)
    creatine = 8.0e-3

    NAA_pred = np.array([NAA_h, NAA_a, NAA_c]) / creatine
    Cho_pred = np.array([Cho_h, Cho_a, Cho_c]) / creatine

    return NAA_pred, Cho_pred
```

Listing 3: Enzyme Kinetics Model - Forward Model (Lines 77–166)

```python
def build_enzyme_model():
    """
    Build PyMC model with enzyme kinetics.

    KEY PARAMETERS:
    - xi_acute, xi_chronic: Noise correlation lengths
    - beta_xi: Protection factor exponent (expect ~2)
    - gamma_coh: Coherence coupling exponent
    - viral_damage_*: Direct damage to enzymes
    - membrane_*: Membrane turnover rates
    """

    with pm.Model() as model:

        # PRIORS
        # Noise correlation lengths (informed by v3.6)
        xi_acute = pm.TruncatedNormal(
            'xi_acute',
            mu=0.50e-9,
            sigma=0.15e-9,
            lower=0.35e-9,
            upper=0.70e-9
        )

        xi_chronic = pm.TruncatedNormal(
            'xi_chronic',
            mu=0.78e-9,
            sigma=0.10e-9,
            lower=0.70e-9,
            upper=0.90e-9
        )

        # Protection factor exponent (informed by v3.6: beta = 1.731)
        beta_xi = pm.TruncatedNormal(
            'beta_xi',
            mu=1.75,
            sigma=0.50,
            lower=0.5,
            upper=3.5
        )

        # Coherence coupling exponent
        gamma_coh = pm.TruncatedNormal(
            'gamma_coh',
            mu=1.5,
            sigma=0.50,
            lower=0.5,
            upper=3.0
        )

        # Viral damage factors (0-1, 1 = no damage)
        viral_damage_acute = pm.Beta('viral_damage_acute', alpha=19, beta=1)
        viral_damage_chronic = pm.Beta('viral_damage_chronic', alpha=9, beta=1)

        # Membrane turnover
        membrane_acute = pm.TruncatedNormal('membrane_acute', mu=2.0, sigma=0.5,
                                            lower=1.0, upper=4.0)
        membrane_chronic = pm.TruncatedNormal('membrane_chronic', mu=1.2, sigma=0.3,
                                              lower=1.0, upper=2.0)

        # FORWARD MODEL
        NAA_pred, Cho_pred = forward_model_enzyme_op(
```

```
            xi_acute, xi_chronic, beta_xi, gamma_coh,
            viral_damage_acute, viral_damage_chronic,
            membrane_acute, membrane_chronic
        )

        # LIKELIHOOD
        sigma_NAA = pm.HalfNormal('sigma_NAA', sigma=0.06)
        sigma_Cho = pm.HalfNormal('sigma_Cho', sigma=0.03)

        NAA_likelihood = pm.Normal('NAA_obs', mu=NAA_pred, sigma=sigma_NAA,
                                   observed=NAA_OBS)
        Cho_likelihood = pm.Normal('Cho_obs', mu=Cho_pred, sigma=sigma_Cho,
                                   observed=CHO_OBS)

        # DERIVED QUANTITIES
        Pi_acute = pm.Deterministic('Pi_acute', (0.8e-9 / xi_acute) ** beta_xi)
        Pi_chronic = pm.Deterministic('Pi_chronic', (0.8e-9 / xi_chronic) ** beta_xi)
        protection_ratio = pm.Deterministic('protection_ratio', Pi_acute / Pi_chronic)

    return model
```

Listing 4: Enzyme Kinetics Model - PyMC Model (Lines 172–314)

##### 6.3 Statistical Improvements Module

**File:** `scripts/statistical_improvements.py`

This module implements enhanced statistical reporting including Bayes Factors, effect sizes, and meta-analysis.

```
def calculate_bayes_factor_savage_dickey(
    posterior_mean: float,
    posterior_sd: float,
    prior_mean: float,
    prior_sd: float,
    null_value: float = 0.0
) -> BayesFactorResult:
    """
    Calculate Bayes Factor using Savage-Dickey density ratio.

    BF_10 = P(theta=theta_0 | prior) / P(theta=theta_0 | posterior)

    For testing H0: xi_acute >= xi_chronic (no protection)
    vs H1: xi_acute < xi_chronic (protective effect)
    """
    # Prior density at null
    prior_density = stats.norm.pdf(null_value, prior_mean, prior_sd)

    # Posterior density at null
    posterior_density = stats.norm.pdf(null_value, posterior_mean, posterior_sd)

    # Avoid division by zero
    if posterior_density < 1e-300:
        bf_10 = np.inf
        log_bf_10 = np.inf
    else:
        bf_10 = prior_density / posterior_density
        log_bf_10 = np.log(prior_density) - np.log(posterior_density)

    # Interpretation (Kass & Raftery 1995)
    if bf_10 > 100:
        interpretation = "Decisive evidence for H1"
    elif bf_10 > 30:
        interpretation = "Very strong evidence for H1"
    elif bf_10 > 10:
```

```
        interpretation = "Strong evidence for H1"
    elif bf_10 > 3:
        interpretation = "Substantial evidence for H1"
    elif bf_10 > 1:
        interpretation = "Weak evidence for H1"
    else:
        interpretation = "Evidence favors H0"

    return BayesFactorResult(
        bf_10=bf_10,
        log_bf_10=log_bf_10,
        interpretation=interpretation,
        h0_description="xi_acute >= xi_chronic (no protective effect)",
        h1_description="xi_acute < xi_chronic (protective paradox)",
        method="Savage-Dickey density ratio"
    )
```

Listing 5: Statistical Improvements - Bayes Factor (Lines 145–196)

```
def random_effects_meta_analysis(
    effects: np.ndarray,
    variances: np.ndarray,
    method: str = 'DL'
) -> MetaAnalysisResult:
    """
    Perform random-effects meta-analysis using DerSimonian-Laird method.

    Parameters
    ----------
    effects : array
        Effect sizes from each study
    variances : array
        Variance of effect sizes
    method : str
        'DL' for DerSimonian-Laird, 'REML' for restricted maximum likelihood

    Returns
    -------
    MetaAnalysisResult with pooled effect, heterogeneity statistics
    """
    k = len(effects)

    if k < 2:
        return MetaAnalysisResult(
            pooled_effect=effects[0] if k == 1 else np.nan,
            pooled_se=np.sqrt(variances[0]) if k == 1 else np.nan,
            pooled_ci=(np.nan, np.nan),
            q_statistic=np.nan,
            q_pvalue=np.nan,
            i_squared=0.0,
            tau_squared=0.0,
            n_studies=k,
            method=method
        )

    # Fixed-effects weights
    w = 1 / variances

    # Fixed-effects pooled estimate
    theta_fe = np.sum(w * effects) / np.sum(w)

    # Q statistic (heterogeneity test)
    Q = np.sum(w * (effects - theta_fe)**2)
    df = k - 1
    q_pvalue = 1 - stats.chi2.cdf(Q, df)

```

```python
    # DerSimonian-Laird estimate of tau^2
    c = np.sum(w) - np.sum(w**2) / np.sum(w)
    tau2 = max(0, (Q - df) / c)

    # Random-effects weights
    w_re = 1 / (variances + tau2)

    # Random-effects pooled estimate
    theta_re = np.sum(w_re * effects) / np.sum(w_re)
    se_re = np.sqrt(1 / np.sum(w_re))

    # 95% CI
    ci = (theta_re - 1.96*se_re, theta_re + 1.96*se_re)

    # I^2 statistic
    if Q > 0:
        i_squared = max(0, ((Q - df) / Q) * 100)
    else:
        i_squared = 0.0

    return MetaAnalysisResult(
        pooled_effect=theta_re,
        pooled_se=se_re,
        pooled_ci=ci,
        q_statistic=Q,
        q_pvalue=q_pvalue,
        i_squared=i_squared,
        tau_squared=tau2,
        n_studies=k,
        method=method
    )
```

Listing 6: Statistical Improvements - Meta-Analysis (Lines 334–411)

#### 6.4 Data Loader Utilities

**File:** quantum/data_loaders.py

This module handles data loading with ART-era classification and mechanistic parameter derivation.

```python
"""
Data loaders for ratio-comparison datasets with ART-era classification and
mechanistic Option A (xi -> Pi_xi) derivation.
"""
from __future__ import annotations

from pathlib import Path
from typing import Literal, Tuple

import numpy as np
import pandas as pd

RATIO_DIR = Path('data/extracted_expanded/data_ratios_comparison')

Era = Literal['pre_modern', 'post_modern', 'unknown']

PAPER_YEAR: dict[str, int] = {
    'Valcour_2015': 2015,
    'Young_2014': 2014,
    'Sailasuta_2012': 2012,
    'Chang_2002': 2002,
}

ART_CUTOFF_YEAR = 2006  # <= pre_modern, >=2007 post_modern
```

```python

def classify_era(year: float | int | None, cutoff: int = ART_CUTOFF_YEAR) -> Era:
    if pd.isna(year):
        return 'unknown'
    try:
        y = int(year)
    except Exception:
        return 'unknown'
    return 'pre_modern' if y <= cutoff else 'post_modern'

def attach_era(df: pd.DataFrame, cutoff: int = ART_CUTOFF_YEAR) -> pd.DataFrame:
    df = df.copy()
    year_series = None
    for col in ['measurement_year', 'scan_year', 'publication_year']:
        if col in df.columns:
            year_series = df[col]
            break

    if year_series is None:
        if 'study' in df.columns:
            df['publication_year'] = df['study'].map(PAPER_YEAR)
            year_series = df['publication_year']
        else:
            df['publication_year'] = np.nan
            year_series = df['publication_year']

    df['art_era'] = [classify_era(y, cutoff) for y in year_series]
    df['art_era_idx'] = df['art_era'].map({'pre_modern': 0, 'post_modern': 1, 'unknown':
    -1})
    return df

def add_derived_covariates(df: pd.DataFrame) -> pd.DataFrame:
    """Compute derived covariates when inputs exist."""
    df = df.copy()
    if 'pVL' in df.columns:
        with np.errstate(divide='ignore'):
            df['log_VL'] = np.log10(pd.to_numeric(df['pVL'], errors='coerce').clip(lower
    =1.0))
    if {'CD4', 'CD8'}.issubset(df.columns):
        with np.errstate(divide='ignore', invalid='ignore'):
            c4 = pd.to_numeric(df['CD4'], errors='coerce')
            c8 = pd.to_numeric(df['CD8'], errors='coerce')
            df['CD4_CD8_ratio'] = c4 / c8
    return df

def compute_Pi_xi_from_xi_nm(xi_nm: pd.Series, beta_xi: float = 1.89,
                             xi_baseline_nm: float = 0.66) -> pd.Series:
    xi = pd.to_numeric(xi_nm, errors='coerce').astype(float).clip(lower=1e-9)
    return (xi / float(xi_baseline_nm)) ** (-beta_xi)

def load_enzyme_inputs(ratio: str, beta_xi: float = 1.89,
                       xi_baseline_nm: float = 0.66) -> Tuple[pd.DataFrame, Path]:
    path = RATIO_DIR / f'enzyme_inputs_{ratio}.csv'
    if not path.exists():
        raise FileNotFoundError(f"Missing enzyme inputs for ratio '{ratio}': {path}")
    df = pd.read_csv(path)
    df = attach_era(df)
    if 'xi_estimate_nm' in df.columns:
        df['Pi_xi'] = compute_Pi_xi_from_xi_nm(df['xi_estimate_nm'],
                                               beta_xi=beta_xi,
                                               xi_baseline_nm=xi_baseline_nm)
```

```
        df['computed_Pi_xi'] = df['Pi_xi']
    else:
        raise ValueError("enzyme_inputs CSV must include 'xi_estimate_nm' column")
    df = add_derived_covariates(df)
    return df, path

def load_bayesian_inputs(ratio: str) -> Tuple[pd.DataFrame, Path]:
    path = RATIO_DIR / f'bayesian_inputs_{ratio}.csv'
    if not path.exists():
        raise FileNotFoundError(f"Missing bayesian inputs for ratio '{ratio}': {path}")
    df = pd.read_csv(path)
    df = attach_era(df)
    df = add_derived_covariates(df)
    return df, path
```

Listing 7: Data Loaders (Complete File)

#### 6.5 Software Environment

Table 10: Software dependencies and versions used in this analysis.

| Package | Purpose | Version |
|---|---|---|
| Python | Programming language | 3.11+ |
| PyMC | Bayesian inference | 5.x |
| ArviZ | Posterior analysis | 0.18+ |
| NumPy | Numerical computing | 1.24+ |
| Pandas | Data manipulation | 2.0+ |
| SciPy | Statistical functions | 1.11+ |
| Matplotlib | Visualization | 3.8+ |
| scikit-learn | Machine learning utilities | 1.5.0+ |
| tqdm | Progress bars | 4.66.3+ |

#### 6.6 Reproducibility

All analysis code, data, and results are archived at Zenodo (DOI: 10.5281/zenodo.18685010) and maintained at `https://github.com/Nyx-Dynamics/noise_decorrelation_hiv`.

To reproduce the complete analysis pipeline:

1. Clone the repository:
```
git clone https://github.com/Nyx-Dynamics/noise_decorrelation_hiv.git
cd noise_decorrelation_hiv
```

2. Create a virtual environment and install dependencies:
```
python3 -m venv .venv && source .venv/bin/activate
pip install -r requirements.txt
```

3. Reproduce all results (models v3.6, v1.0, v2.0, v4.0):
```
python reproduce_all.py
```
or equivalently: `make reproduce`

All random seeds are fixed for deterministic reproduction. The pipeline runs four convergent Bayesian models, generates all manuscript figures, and produces cryptographic checksums for output verification. Expected runtime on a modern workstation (8+ cores, 16 GB RAM): approximately 30–60 minutes for full MCMC sampling across all models. Individual models can also be run independently (see `QUICKSTART.md`).
